# STIP1/HOP Promotes the Formation of Cytotoxic α-Synuclein Oligomers

**DOI:** 10.1101/2025.03.26.645247

**Authors:** Benjamin S. Rutledge, Carter J. Wilson, Rachel M. Lau, Juan C. Jurado-Coronel, Esther del Cid-Pellitero, Mikko Karttunen, Thomas M. Durcan, Edward A. Fon, Marco A.M. Prado, Justin Legleiter, Martin L. Duennwald, Wing-Yiu Choy

**Author notes:** To whom correspondence may be addressed: Dr. Wing-Yiu Choy, Dr. Martin Duennwald, Dr. Justin Legleiter.

## Abstract

The accumulation of alpha-synuclein (a-Syn) as toxic oligomers, and subsequently in Lewy bodies, is a pathological hallmark of Parkinson’s disease (PD) and other synucleinopathies. Molecular chaperones and co-chaperones are expected to act in concert to maintain physiological activities of proteins, including a-Syn, but in neurodegeneration this process can become mal-adaptive. Transcript levels of Stress inducible phosphoprotein 1 (STIP1), a co-chaperone of Hsp90/Hsp70, are elevated in brain samples from PD patients. In synucleinopathy mouse models, STIP1 has unexpected bidirectional effects on a-Syn, with overexpression of STIP1 aggravating a-Syn toxicity, whereas knockdown of STIP1 improves toxicity and behavioural phenotypes. However, it is unclear how STIP1 enhances the toxicity of a-Syn. Here we unravel the mechanisms by which the direct interaction between STIP1/HOP and a-Syn regulates the neurotoxicity of a-Syn. Specifically, two binding motifs in the C-terminus of a-Syn directly interact with the TPR2A domain of STIP1/HOP in a dynamic manner, competing for a shared interface on TPR2A. Binding of STIP1/HOP to a-Syn attenuates the formation of a-Syn fibrils while promoting the accumulation of high molecular weight amorphous a-Syn species. Samples of a-Syn aggregated in the presence of STIP1/HOP contain significantly more A11-positive oligomeric species and cause a greater reduction in cell viability than a-Syn aggregated in the absence of STIP1/HOP in neuronal cells. Our results provide a mechanism by which the direct interaction between STIP1/HOP and the C-terminus of a-Syn promotes the formation of cytotoxic, non-amyloidogenic, high molecular weight a-Syn species. Our model offers an explanation for the unexpected pathological link between STIP1 and a-Syn toxicity, thus opening new therapeutic avenues for the treatment of synucleinopathies.

## Introduction

The aggregation of alpha-synuclein (a-Syn), a presynaptic neuronal protein, and its deposition in Lewy bodies is a hallmark of Lewy Body Dementia (LBD), Parkinson’s disease (PD) and other synucleinopathies [1–3]. a-Syn consists of three domains: an amphipathic N-terminus, the aggregation-prone central non-Aβ component (NAC), and an acidic C-terminus [4]. Due to its lack of stable structural elements, a-Syn can be categorized as an intrinsically disordered protein (IDP), which samples an ensemble of diverse conformations. Under pathological conditions, a-Syn is susceptible to adopting conformations that expose its NAC domain, facilitating the formation of toxic polymorphic oligomers and, subsequently, intracellular inclusions of fibrillar a-Syn [5]. Although aggregation has previously been attributed to the NAC region, recent studies indicate that the terminal domains of a-Syn can mediate oligomerization, aggregation, and pathology [6–10]. Accordingly, the N-terminal domain and adjacent residues of a-Syn are central for the neurotoxicity associated with a-Syn oligomers and contain the majority of PD-associated mutations, such as A18T, A29S, A30P, E46K, G51D, A53T, and E83Q [6,11–16]. The C-terminal domain, on the other hand, contributes substantially to the energetic barriers that regulate the transition between monomeric, oligomeric, and fibrillar conformers of a-Syn [17–19].

Numerous cellular protein quality control (proteostasis) pathways, such as molecular chaperones and the ubiquitin-proteosome system, participate in modulating a-Syn accumulation and toxicity [20–22]. For instance, up to 90% of a-Syn in live cells is estimated to be engaged in interactions with molecular chaperones [23]. The N-terminus of a-Syn, together with a hydrophobic region around Tyr39, dictates the specificity of the interactions between a-Syn and multiple molecular chaperones [23]. Chaperone binding modulates a-Syn aggregation often through an ATP-independent mechanism, as opposed to the ATP-dependent folding of many other client proteins [24]. Two major cytosolic chaperones, Hsp90 and Hsp70, interact with and shield hydrophobic residues in the N-terminal and NAC regions exposed in non-native aggregation-prone conformers of a-Syn [25–29], thereby inhibiting a-Syn fibrillization and favouring monomeric non-toxic soluble a-Syn conformers *in vitro* [26–28,30]. By contrast, only few interactions mediated by the C-terminal domain of a-Syn have been described, including interactions with calcium [31], tau [32], fatty acid binding protein 3 (FABP3) [33], and the prion protein (PrP^C^) [34,35]. As opposed to N-terminal interactions with chaperones such as Hsp90 and Hsp70, these described C-terminal interactions appear to promote a-Syn oligomerization and enhance a-Syn toxicity [31–33,36,37].

Stress inducible phosphoprotein 1 (STIP1), also known as Hsp-organizing protein (HOP) in humans, is a major co-chaperone to both Hsp90 and Hsp70, facilitating the transfer of client proteins between these two chaperones and regulating their ATPase activity [38–40]. STIP1 contains three tetratricopeptide repeat (TPR) domains (TPR1, TPR2A, and TPR2B) and two aspartate and proline rich (DP) domains (DP1 and DP2). STIP1 interacts with Hsp90 via its TPR2A domain and with Hsp70 through its TPR1 and TPR2B domains [39–42]. Although largely known for its role as a Hsp70 and Hsp90 co-chaperone, STIP1 also regulates other proteostatic functions, including proteasomal degradation [43,44], actin dynamics [45], and support of kinase activity [46]. STIP1 can also be secreted into the extracellular space [47,48], where it functions in cell signaling [49].

STIP1 has complex effects on several neurodegeneration-associated proteins, including TAR DNA-binding protein 43 (TDP-43) [50], PrP^C^ [51], amyloid-β [52], and a-Syn [36]. Transcript levels of STIP1 are elevated in brain samples from PD patients as compared to healthy individuals [36]. Experiments in a synucleinopathy mouse model indicate that increased levels of STIP1 exacerbate a-Syn aggregation, increase pS129 a-Syn (phosphorylated at serine 129) levels, and enhance neurotoxicity. Conversely, decreased levels of STIP1 partially inhibit a-Syn aggregation, reduce the levels of pS129 a-Syn, and mitigate behavioral defects [36], suggesting a dose-dependent relationship between STIP1 function and a-Syn toxicity. STIP1 associates with human a-Syn in mouse brains, with the TPR2A domain of STIP1 interacting directly with the C-terminus of a-Syn [36]. However, the mechanism by which STIP1 modulates a-Syn toxicity are still enigmatic.

In the present work, we reveal the dynamic interaction between the TPR2A domain of STIP1/HOP and two motifs of negatively charged residues in the C-terminus of a-Syn. We find that this interaction stabilizes soluble, amorphous non-fibrillar conformers of a-Syn, while diminishing the formation of larger a-Syn fibrils. Remarkably, the amorphous non-fibrillar species resemble pathological a-Syn oligomers and are highly toxic to cells. Our study uncovers the molecular mechanisms by which STIP1/HOP interactions favour a toxic oligomeric state of a-Syn. We propose an unexpected pathological link between detrimental proteostasis functions of STIP1 and a-Syn toxicity as a mechanism that can be targeted in synucleinopathies.

## Materials and Methods

Peptides derived from the amino acid sequence of a-Syn (a-Syn 110-140: Ac-EGILEDMPVDPDNEAYEMPSEEGYQDYEPEA-COOH; Biotin – a-Syn 110-140: Biotin-EGILEDMPVDPDNEAYEMPSEEGYQDYEPEA-COOH; pS129 a-Syn 110-140: Ac-EGILEDMPVDPDNEAYEMP[pS]EEGYQDYEPEA-COOH; S1: Ac-PVDPDNEAYEM-NH_2_, S2: Ac-EEGYQDYEPEA-COOH) and the Hsp90 pentapeptide (Ac-MEEVD-COOH) were synthesized by Tufts University Core Facility. Peptides were dissolved in 40% 20 mM HEPES and 50 mM NaCl at pH 7.2 and 60% 0.1 M ammonium bicarbonate. After dissolving, peptides were dialysed using 1 mL 100-500 Da MWCO dialysis cassettes into 20 mM HEPES and 50 mM NaCl at pH 7.2. Final concentrations were confirmed by amino acid analysis (SPARC BioCentre, The Hospital for Sick Children, Toronto).

### Recombinant Protein Purification

Human wild type a-Syn and a-Syn mutants (A53T, E137A, D119A/D121A, D135A/E137A/E139A, and D119A/D121A/D135A/E137A/E139A) generated using QuikChange Multi Site-Directed Mutagenesis (Agilent Technologies) were expressed in *E. coli* BL21 (DE3) cells in minimal M9 media containing either ^14^NH_4_Cl (1 g/L) or ^15^NH_4_Cl (1 g/L) as the sole nitrogen source. Recombinant human a-Syn and mutational variant expression and purification was performed as previously described [36]. All NMR and aggregation kinetics studies were conducted immediately following the purification of a-Syn to avoid any potential influence of storage at −80 °C.

pDEST17 (Invitrogen, Carlsbad, CA, USA) expression plasmids encoding full-length mouse STIP1, TPR1 domain (mTPR1; residues 1–118), TPR2A domain (mTPR2A; residues 217–352), and the TPR2A domain of human HOP (hTPR2A; residues 217–352) fused with N-terminal tobacco etch virus (TEV) cleavable hexa-His tags were transformed into BL21 (DE3) pLysS for protein expression in minimal M9 media containing either ^14^NH_4_Cl (1 g/L) or ^15^NH_4_Cl (1 g/L) as the sole nitrogen source. Expression and purification of full-length STIP1/HOP and the TPR domains were performed as previously described [53].

Protein concentrations were determined by measuring absorbance at 280 nm using a NanoDrop spectrophotometer. For all samples used for determining binding kinetics, protein concentrations were further confirmed by amino acid analysis (SPARC BioCentre, The Hospital for Sick Children, Toronto).

### S129 Phosphorylation of α-synuclein using PLK3

Phosphorylation of a-Syn and A53T a-Syn at S129 was done as previously described [36]. Reactions containing approximately 5 mg of a-Syn, 4 µg of Polo-like kinase 3 (PLK3) (ThermoFisher PV3812), 50 mM HEPES, 10 mM MgCl_2_, 1 mM EGTA, 1 mM DTT, and 2 mM ATP were incubated overnight at 30 °C. Following incubation, samples were centrifuged at 16,000 x g for 15 minutes at 4 °C to remove any aggregates formed during incubation. Samples were exchanged by dialysis overnight using a 5 mL 500-1000 Da MWCO dialysis cassette with 20 mM HEPES and 50 mM NaCl at pH 7.2. The protein was concentrated with a 5 kDa MWCO Vivaspin Turbo 4 centrifugal concentrator (Sartorius) to a volume of 1 mL and Lowry protein assay was used to estimate the concentration. Concentrations were later confirmed by amino acid analysis (SPARC BioCentre, The Hospital for Sick Children, Toronto).

### Solution NMR spectroscopy

NMR experiments were performed on a 600 MHz Varian Inova/Bruker spectrometer equipped with an xyz-gradient triple resonance probe. The ^1^H-^15^N HSQC spectra were collected using 128 × 537 complex points in the ^15^N and ^1^H dimensions, respectively. Experiments utilizing ^15^N labeled a-Syn variants were conducted at 10 °C in 20 mM HEPES and 50 mM NaCl at pH 7.2. For experiments utilizing ^15^N labeled TPR domains, all experiments were conducted at 25 °C in 20 mM HEPES and 50 mM NaCl at pH 7.2. NMR data were processed using NMRPipe [54] and analyzed using NMRViewJ [55] software. Chemical shift analyses of titration sets were conducted using the Titration Analysis function in NMRViewJ. Chemical shift perturbations for ^1^H-^15^N peaks were calculated following Δδ = [(Δδ_1H_)^2^ + (0.14*Δδ_15N_)^2^]^1/2^, where Δδ_1H_ and Δδ_15N_ are the changes in ^1^H and ^15^N chemical shifts (in ppm) upon the addition of binding target [56].

### Molecular Dynamics Simulations

Initial configurations comprised residues 111-140 of human a-Syn (N-capped with acetyl group) and human TPR2A. 24 semi-extended configurations of α-synuclein were uniformly positioned around TPR2A (PDB: 1ELR). In each case the distance between at least a pair of heavy atoms (one from each protein) was within 1 nm. These configurations were centered in a 10 × 10 × 10 nm cubic box and solvated using mTIP3P [57] (the prefix m denotes the CHARMM36 [58] compatible TIP3P water model) or TIP4P-D water [59]. K^+^ and Cl^−^ counterions were added to ensure neutrality.

All simulations were performed using the GROMACS [60] software and the CHARMM36m force field [58]. An initial energy minimization was performed using the steepest descents algorithm. The temperature was maintained at 298.15 K using the Parrinello-Donadio-Bussi velocity rescaling method [61] with a coupling time of 1.0 ps, while the pressure was maintained at 1 bar using the Parrinello-Rahman barostat [62] with a coupling time constant of 5.0 ps. The simulation time step was 2.0 fs. The temperature and pressure were chosen to match physiological conditions. Long-range electrostatic interactions were calculated using the Particle-mesh Ewald (PME) method [63] with a Fourier spacing of 0.12 nm and a real-space cut-off of 1.0 nm. Lennard-Jones interactions were force-switched off between 1 nm and 1.2 nm.

Bond lengths were constrained using the Parallel LINear Constraint Solver [64]. Simulations were run for 1.8 µs each, this gave 24 simulations with mTIP3P and 24 simulations with TIP4P-D; the latter was included because it produces highly extended disordered ensembles that have been suggested to well represent experimental observations. These were combined into one large trajectory with frames stored every 1 ns.

### Biolayer Interferometry

An Octet R2 biolayer interferometry instrument (Sartorius) equipped with the Octet BLI Discovery v12.2 and Octet Analysis Studio v12.2 software was used to measure the binding affinities of the a-Syn 110-140, S1, S2, and Hsp90 pentapeptide for hTPR2A. Biotinylated hTPR2A was produced by incubating 15 µM hTPR2A with Biotin-maleimide (Sigma B1267) in 20 mM HEPES and 50 mM NaCl at pH 7.2 for 30 minutes at room temperature. After incubation, the sample was buffer exchanged into fresh 20 mM HEPES and 50 mM NaCl at pH 7.2 using a 5 kDa MWCO Vivaspin Turbo 4 (Sartorius) concentrator with 4 sample volumes of fresh buffer to remove unreacted biotin-maleimide. Biotinylated hTPR2A or the Biotin-a-Syn 110-140 peptide were immobilized onto streptavidin (SA) biosensors (Satorius) by incubating the sensor in 3 µM of biotinylated hTPR2A or the Biotin-a-Syn 110-140 peptide for 10 minutes. The biosensors were then quenched in 1 mM free biotin for 10 minutes to prevent non-specific binding. The biosensors were then dipped into wells of increasing concentrations of ligand for 60-120 seconds and dissociation was monitored by dipping the biosensor in 20 mM HEPES and 50 mM NaCl at pH 7.2 supplemented with 0.1% BSA and 0.02% Tween 20.

The assays were performed using 96 well plates at 25 °C with 1,000 rpm agitation in 20 mM HEPES and 50 mM NaCl at pH 7.2 supplemented with 0.1% BSA and 0.02% Tween 20 to prevent non-specific binding. An unloaded SA biosensor was dipped into buffer alone and the subsequent ligand samples as a control. The Biotin-a-Syn 110-140 peptide was immobilized and the binding of hTPR2A was fit to a heterogeneous binding model to calculate the binding affinity. Biotinylated hTPR2A was immobilized to monitor the binding of the Hsp90 pentapeptide as well as the a-Syn C-terminal, S1, and S2 a-Syn peptides, all of which were fit to a one-to-one binding model to determine binding affinity.

### ThT aggregation assays

For all aggregation experiments, wild type a-Syn was used unless stated otherwise. Thioflavin T (ThT) dye was used at a final concentration of 10 µM to monitor the aggregation of a-Syn (concentrations ranging from 20 µM to 60 µM based on experimental design) using the excitation wavelength of 445 nm and emission wavelength of 485 nm. Samples contained wild type or variant a-Syn with the absence or presence of various concentrations of the TPR domains and in the absence or presence of peptides. The reaction buffer consisted of phosphate buffer saline (PBS) at pH 7.4 with 0.1% sodium azide to prevent bacterial growth. For delayed addition of hTPR2A, concentrated hTPR2A (>1.5 mM) was added at equimolar concentration to a-Syn directly into the well for each sample at its corresponding time point to avoid significant volume changes. Each experiment was performed with 4 replicates and values were averaged. Each replicate consisted of a 50 µL final reaction volume and replicates were monitored in black non-binding half volume 96-well plates with clear bottoms (Corning Incorporated). A sterile glass bead (∼2.7 mm diameter) was added to each well to aid with sample mixing. A Cytation 5 Cell Imaging Multi-Mode Reader (Biotek, Winnoski, VT, USA) was used at a temperature of 37 °C with continuous orbital shaking at 425 rpm.

### Electron Microscopy

Fibrillar a-Syn samples were produced by aggregating 175 µM a-Syn in the absence or presence of equimolar mTPR1 or hTPR2A for 5 days at 37 °C in a shaking incubator (400 rpm). Samples were characterized using a negative staining. They were added to 200 mesh carbon coated copper grid (3520C-FA, SPI Supplies), fixed with 4% PFA and stained with 2% acetate uranyl (22400-2, EMS) for 1 minute. Aggregated a-Syn samples were visualized using a transmission electron microscope (Tecnai G2 Spirit Twin 120 kV Cryo-TEM) coupled to a camera (Gatan Ultrascan 4000 4 k × 4 k CCD Camera System Model 895) and analyzed with Fiji-ImageJ1.5 and GraphPad Prism 9 software.

### Size exclusion chromatography

a-Syn at a concentration of 300 µM was incubated in the absence and presence of hTPR2A at an equimolar concentration in 20 mM HEPES and 50 mM NaCl at 37 °C with a shaking speed of 400 rpm. A sterile glass bead (∼2.7 mm diameter) was added to each sample to aid with mixing. To ensure that hTPR2A did not form high molecular weight species itself, a control sample of 300 µM hTPR2A was subjected to the same aggregation conditions. All samples were run on a Superdex 200 Increase 10/300 GL (Cytiva) column with a flow rate of 0.6 mL/min. Elution was collected and analyzed by SDS-PAGE using Coomassie Blue stain.

### Atomic Force Microscopy

The a-Syn, mTPR1, and hTPR2A mixtures were prepared in the same manner as for other ThT assays. To remove any possible contaminants, protein stocks were filtered using a 0.8 µM syringe filter before preparing the mixtures. Incubations of a-Syn alone and with equimolar amounts of mTPR1 or hTPR2A were shaken at 400 rpm on an orbital shaker, and each incubation contained a sterile glass bead (∼2.7 mm diameter). At various time points, 2 µL aliquots of each condition were taken and deposited on freshly cleaved mica, allowed to sit on the substrate for 1 min, were then washed with 100 μL of 18 MΩ water, and dried with a gentle stream of nitrogen. Imaging of samples was performed on a Nanoscope V Multimode Atomic Force Microscope (AFM) (Veeco, Santa Barbara, CA) with a closed-loop vertical engage J-scanner. All images were obtained in tapping mode utilizing a diving-board-shaped silicon oxide cantilever with a spring constant of 40 N/m and the resonance frequency of ∼300 kHz.

AFM image processing and analysis was performed in Matlab (MathWorks, Natick, MA) by utilizing the imaging processing toolbox as previously described [65]. A 2^nd^ order polynomial flattening algorithm was used to correct for background curvature. A height threshold (z = 0.9 nm) was implemented to produce binary maps used to identify discreet aggregates in each image. Physical features of individual aggregates (e.g., height, volume, diameter, and area covered) were measured. Each aggregate was also assigned a tracking number. A minimum of five 5 × 5 µm AFM images for each time point was analyzed, resulting in a minimum of 125 µm^2^ of surface area analyzed. As fibrils and fibril bundles are typically larger and demonstrate an elongated morphology relative to oligomers and amorphous aggregates, data sets were filtered for a minimum area occupied by the aggregate (75 pixels) and aspect ratio (> 2) to identify fibrils and determine fibril load. The reliability of these filters to correctly identify fibrils was validated by labeling the aggregates that passed the filters directly in the images. Aggregates that did not pass the morphological filters were categorized as oligomers or amorphous aggregates.

### Dynamic Light Scattering

Samples from a ThT assay conducted using 20 µM a-Syn in the absence and presence of equimolar hTPR2A were collected after 3 and 6 days of aggregation. Samples were pulse spun briefly before dynamic light scattering measurements. The upper portion of the sample was used to avoid dust contamination from the sample but maintain high molecular weight species and fibrils. Measurements were done using a Dynapro NanoStar dynamic light scattering instrument (Wyatt Technologies) with an incident light wavelength of 825 nm and a scattering angle of 90°. For each sample, 20 consecutive autocorrelation functions, each with a 5 second accumulation time were acquired and the autocorrelation functions of the 10 lowest scattering acquisitions were averaged for hydrodynamic size calculations. Deconvolution was performed using the accompanying DYNAMICS software (v7.1.9) between 1.5 μs and 100 ms for all averaged curves using identical resolution settings for each sample. All calculations used a solvent index of 1.333 and a viscosity of 1.019 g m^−1^ s^−1^.

### Dot Blot Assay

The a-Syn (70 μM) aggregation reactions in PBS buffer were incubated at 37 °C in a shaking incubator (500 rpm) in the presence or absence of equimolar hTPR2A for 1, 2, 3, 5, or 7 days to induce aggregation. Aggregation reactions were run in triplicate for each time point. For each time point, the sample was diluted to a final concentration of 15 μM, 3 μM, and 0.6 μM. A nitrocellulose membrane was pre-wet by soaking it in PBS for 5 minutes before placing it in a Minifold I Dot-Blot 96-Well System (10447900, VWR) and applying vacuum. Each well was washed with 100 μL of PBS using low vacuum. For each dilution of the aggregation samples, 10 μL was blotted on the membrane using low vacuum and the membrane was allowed to dry. The membrane was then blocked using 5% skim milk powder (Carnation/Nestle, Vevey, Switzerland) in Tris-buffered saline with 0.05% (v/v) Tween 20 (Thermo Fisher Scientific, Waltham, MA, USA) (TBST) and incubated for 1 hour on a shaker at room temperature. Following blocking, the membrane was washed with 10 mL aliquots of TBST for 10-minute intervals over a 30-minute period. The membrane was then incubated overnight with primary A11 antibody (Invitrogen, Carlsbad, CA, USA, AHB0052) diluted in TBST with 5% (w/v) Bovine serum albumin (BSA) (Wisent Inc, Quebec, Canada) on a shaking incubator at 4 °C. The membrane was washed again using 10 mL aliquots of TBST at 10-minute intervals for 30 minutes and then incubated in Rabbit secondary antibody conjugated with Alexa680 diluted in TBST with 1% (w/v) BSA on a shaker for 1 hour at room temperature. Following incubation, the membrane was washed in 10 mL of TBST and the membrane was imaged using the ChemiDoc MP Imaging System (Bio-Rad, Hercules, CA, USA) and analyzed using Image Lab (Bio-Rad, Hercules, CA, USA).

### Yeast Strains and Yeast Media

The yeast strain BY 4741 (MAT α his3Δ1 leu2Δ0 lys2Δ0 ura3Δ0) was used in this study. Yeast-peptone-dextrose (YPD)-rich media (10 g/L of yeast extract, 20 g/L of peptone, and 20 g/L of dextrose) and selective dextrose (SD) media (6.7 g/L yeast nitrogen base (YNB), 60 mg/L of L-isoleucine, 20 mg/L of L-arginine, 40 mg/L of L-lysine HCl, 60 mg/L of L-phenylalanine, 10 mg/L of L-threonine, and 2% glucose) in either liquid media or agar plates (20 g/L) were used to grow yeast cells. SD media was supplemented with four different amino acids (40 mg/L of L-tryptophan, 60 mg/L of L-leucine, 20 mg/L of L-histidine-monohydrate, 10 mg/L of L-methionine, and 20 mg/L of uracil) depending on the selective marker of the plasmid. To expose cells with the established Hsp90 inhibitor, Radicicol, liquid media or plates were supplemented with 10 µg/mL of Radicicol.

### Yeast Transformations

Yeast transformations were performed utilizing the standard PEG/lithium acetate transformation method [66].

### Growth Assays (Spotting assays) and Quantification

The relative growth of the yeast cells was measured using spotting assays and quantification as described by Petropavlovskiy et al [67]. In summary, 3 mL of SD media was inoculated with cells and incubated in a shaking incubator at 30 °C overnight. A 48-prong frogger (V&P Scientific, San Diego, CA, USA) was used to spot normalized serial dilutions of cells onto SD plates lacking the selective amino acid. For Radicicol treated plates, Radicicol was added to SD media to a final concentration of 10 µg/mL immediately before media was poured when warm to the touch. Plates were incubated at either 30 °C or 37 °C and growth was monitored during the entire growth period. All spotting assays contain respective control lanes on each experimental plate, and images of the plates were cropped and separated to highlight relative side-by-side comparisons. Unless otherwise indicated, the pixel count of the third dilution of the growth assay was used for quantification. Values were normalized to the average of the respective control lanes. For all growth assays, a minimum of three biological replicates were used.

### Fluorescent Microscopy and Quantification

SD media was inoculated with yeast expressing fluorescently tagged constructs and incubated overnight at either 30 °C or 37 °C in a shaking incubator. For Radicicol treatment, cultures were treated with 10 μg/ml Radicicol 6 hours prior to imaging. Each culture (1-2 μL) was placed on a microscope slide and imaged using a Zeiss Axio Vert.A1 microscope. For each condition, three biological replicates were imaged, and three images of random fields of each biological replicate were imaged. The extent of a-Syn foci formation was quantified as the percentage of cells that contained a-Syn foci as described by Petroi et al [68]. Each data point represents the average percentage of cells that contained foci across the three images taken for each respective biological replicate. For each biological replicate, a minimum of 100 cells was counted towards the percent of cells containing foci unless otherwise stated.

### Mammalian cell culture and media

The SH-SY5Y cell line was used in the course of this study. Cells were grown in Dulbecco’s Modified Eagle Medium (DMEM, Corning, Corning, NY, USA) with 4.5 g/L glucose. Media was supplemented with 10% fetal bovine serum (FBS, Gibco/Thermo Fisher Scientific, Waltham, MA, USA) and 1X penicillin-streptomycin solution (Corning, Corning, NY, USA). Cells were grown at 37 °C with 5% CO_2_.

### Cytotoxicity Assay

The cytotoxicity assay protocol was adapted from the protocol previously described by Daturpalli et al [26]. Cell viability was measured using the CellTiter-Glo 2.0 Luminescent Cell Viability Assay (Promega, Madison, WI, USA), which determines the number of viable cells in the culture by measuring the amount of available ATP. Cells were split into a 96-well plate and grown for 48 hours at 37 °C DMEM. Each well was seeded with 10,000 cells. The a-Syn (70 μM) aggregation reactions in PBS buffer were incubated at 37 °C in a shaking incubator (500 rpm) in the presence or absence of equimolar hTPR2A for 1, 2, 3, 5, or 7 days to induce aggregation. Aggregation reactions were run in triplicate for each time point. The cell media was exchanged with culture medium supplemented with aggregation samples. The final concentration of a-Syn in the cell media was 35 μM (calculated using the initial monomer concentration). Controls were mixed containing monomeric a-Syn, hTPR2A, and PBS buffer using the same respective volumes of media and buffer as the aggregation samples. The SH-SY5Y cells were incubated for 16 hours in the treated media. The CellTiter-Glo 2.0 Luminescent Cell Viability Assay was carried out following the supplier’s instructions and signal was measured using a Cytation 5 Cell Imaging Multi-Mode Reader (Biotek, Winooski, VT, USA).

### Statistical analysis

Statistical analysis of ThT assays, Dot blots, cell viability, growth assays, and percentage of foci assessed by microscopy were performed using the GraphPad Prism X software (GraphPad Software, Boston, MA). To determine statistical significance, unpaired t-tests were used to compare means and standard deviations between relevant controls and experimental data sets (each data set was composed of a minimum of 3 biological repeats). The P values are provided for all conditions where P < 0.05. Bars represent standard errors of the mean.

## Results

### Two linear motifs in the C-terminal domain of a-Syn mediate its interaction with the TPR2A domain of HOP

Mouse STIP1 shares a high sequence identity (97.4%) with its human homologue (HOP), with only three amino acid positions (residues 243, 250, and 269) differing between STIP1-TPR2A domain (mTPR2A; residues 217–352) and the TPR2A domain of human HOP (hTPR2A). NMR ^1^H-^15^N HSQC experiments were used to confirm that hTPR2A interacts with a-Syn in the same manner as mTPR2A [36]. Measuring the changes (perturbations) in the chemical shifts for each assigned residue allows for the identification of the binding interface and insights into the strength of the interaction. Almost identical chemical shift perturbations (CSPs) are observed in C-terminal residues of ^15^N-labeled a-Syn, pS129 a-Syn, A53T a-Syn, and pS129 A53T a-Syn in the presence of mTPR2A and hTPR2A (Figures S1 and S2). Notably, minor CSPs are observed for some residues in the N-terminal domain, likely due to the loss of transient long-range intramolecular interactions between the N- and C-terminal domains of a-Syn upon mTPR2A/hTPR2A binding [69]. These results clearly indicate that, like the mTPR2A domain of STIP1, the hTPR2A of HOP mediates the interaction with the C-terminal domain of wild type and pathological variants of human a-Syn.

Next, to identify key residues in the C-terminal domain of a-Syn that regulate the binding of hTPR2A, we acquired ^1^H-^15^N HSQC spectra of ^15^N-labeled a-Syn in the presence of increasing concentrations of hTPR2A. Quantitative analysis of the NMR titration series reveals that two negatively charged residues in the middle of the C-terminal domain (D119 and D121) and three negatively charged residues close to the C-terminus (D135, E137, and E139) display large CSPs upon binding hTPR2A (Figs. 1A, 1B, and 1E). Group fitting the CSPs of these residues to a 1:1 binding model yields a dissociation constant of 169 ± 67 µM for a-Syn/hTPR2A binding. To further assess the contributions of these two groups of acidic residues to hTPR2A binding, we repeated the NMR titration experiment with a D119A/D121A double mutant of a-Syn (Fig. 1C). Notable CSPs were no longer observed for the residues neighbouring the D119A and D121A mutation sites, suggesting D119 and D121 are crucial for hTPR2A binding.

**Figure 1.**
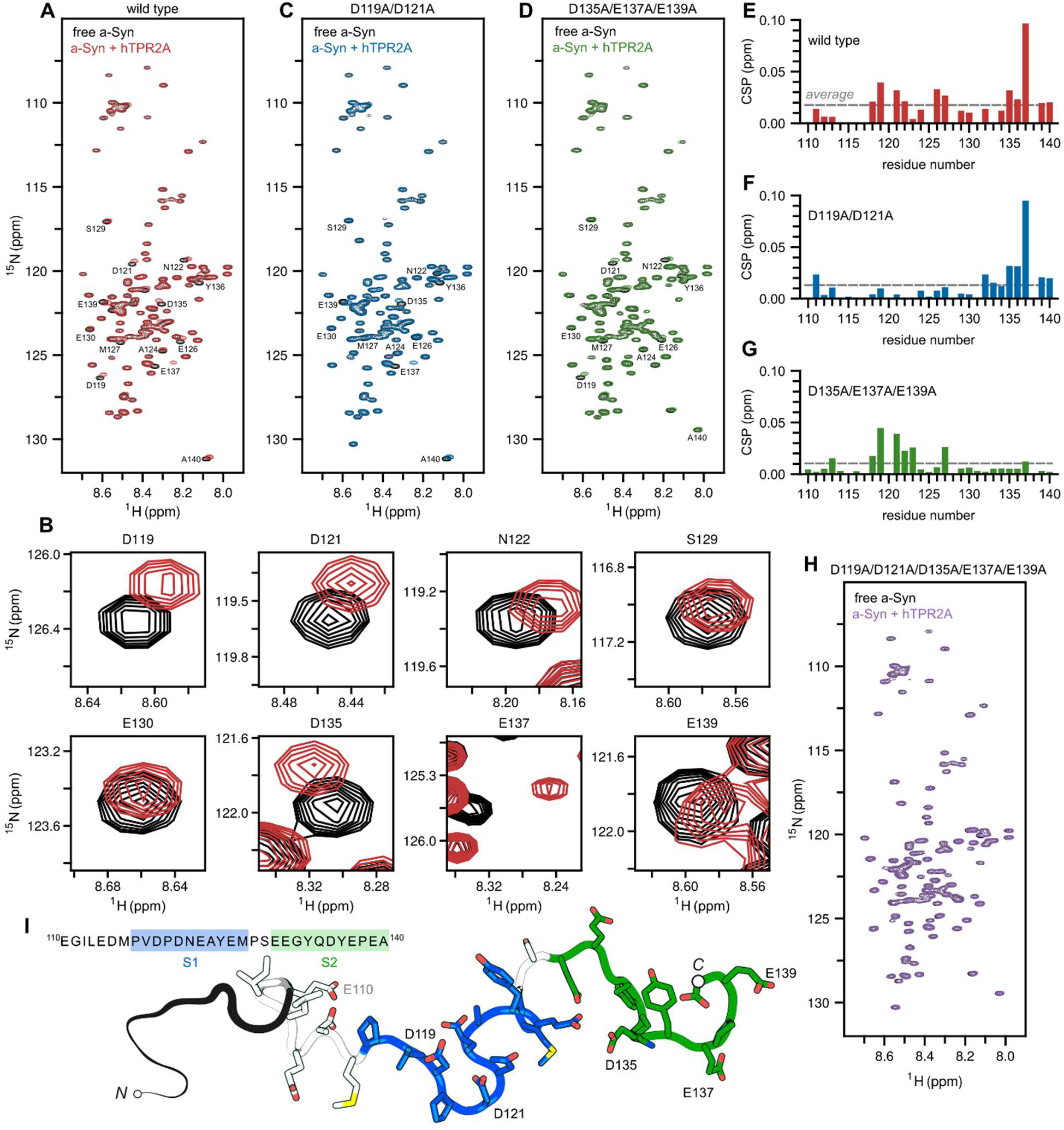
Identification of two motifs in the C-terminal of a-Syn which mediate the interaction with TPR2A. (A) NMR analysis of ^15^N-labeled a-Syn at 100 µM in the absence (black) and the presence of 100 µM hTPR2A (red). (B) Key residues which experience significant chemical shift perturbations upon addition of hTPR2A. (C) ^1^H-^15^N HSQC spectrum of 150 µM D119A/D121A a-Syn in the absence (black) and the presence (blue) of 150 µM of hTPR2A. (D) ^1^H-^15^N HSQC spectrum of 35 µM D135A/E137A/E139A a-Syn in the absence (black) and the presence (green) of 35 µM of hTPR2A. (E-G) Quantification of chemical shift perturbations (CSPs) observed in the C-terminal domain (residues110-140) of a-Syn (E), D119A/D121A a-Syn (F), and D135A/E137A/E139A a-Syn (G) upon addition of equimolar hTPR2A. The average CSP for the C-terminal residues of each respective a-Syn variant is indicated by dashed line. (H) ^1^H-^15^N HSQC spectrum of 35 µM D119A/D121A/D135A/E137A/E139A a-Syn in the absence (black) and the presence (purple) of 35 µM of hTPR2A. (I) Cartoon depiction of the proposed two hTPR2A binding sites in the C-terminal domain of a-Syn, in which site 1 (S1) includes residues D119 and D121, and site 2 (S2) includes residues D135, E137, and E139. Sequences comprising the peptides used for the full C-terminal domain of a-Syn, as well as S1 and S2 individually, are displayed.

Unexpectedly, CSPs observed for D135, E137, and E139 are not substantially reduced, indicating that their interactions with hTPR2A are largely unaffected by the D119A and D121A substitutions (Figs. 1C and 1F). Similarly, D135A, E137A, and E139A substitutions result in no notable CSPs of residues neighbouring the D135A, E137A, and E139A mutation sites upon hTPR2A titration, whereas the CSPs for D119, D121, and neighboring residues were not considerably reduced (Figs. 1D and 1G). Finally, D119A, D121A, D135A, E137A, and E139A substitutions result in no observable CSPs upon the addition of hTPR2A (Fig. 1H). Taken together, these results suggest that the C-terminus of a-Syn contains two motifs that interact independently with hTPR2A, with site 1 (S1) containing the key charged residues D119 and D121 and site 2 (S2) with residues D135, E137 and E139, both of which can bind hTPR2A (Fig. 1I).

### The S1 and S2 motifs compete for binding of overlapping interfaces on hTPR2A

To identify the interfaces on hTPR2A involved in the interaction with the two binding motifs in the C-terminus of a-Syn we performed NMR ^1^H-^15^N HSQC experiments on ^15^N-labeled hTPR2A in the presence of peptides comprising residues 110-140 (C-terminal peptide), residues 117-127 (S1 peptide), and residues 130-140 (S2 peptide) of a-Syn (Figure S3). CSPs were plotted against residue number in hTPR2A and residues experiencing above average CSPs were mapped onto the complex structure of hTPR2A (Figs. 2A-C) [70]. Intriguingly, addition of peptides that cover the C-terminal domain of a-Syn, the S1 motif, and the S2 motif causes CSPs of similar residues within the hTPR2A, indicating that either the S1 or S2 motif alone is sufficient to interact with hTPR2A. Importantly, the results indicate that binding of both S1 and S2 is primarily mediated by the same set of residues in hTPR2A, suggesting that S1 and S2 interact with the same interface on hTPR2A. Notably, the C-terminal peptide does not form additional interactions with hTPR2A beyond S1 and S2 (Figs. 2A-C). We conducted competitive binding NMR experiments to confirm that a-Syn peptides compete with full-length a-Syn for binding hTPR2A. We monitored CSPs of ^15^N-labeled a-Syn in the presence of equimolar hTPR2A upon the addition of C-terminal, S1, or S2 peptide. If the peptides compete with regions of a-Syn for the same binding interface on hTPR2A, the CSPs for those regions of a-Syn are expected to reduce as a larger proportion exists in the unbound state. The addition of equimolar S1 or S2 peptide reduces the CSPs of key residues within both S1 (D119 and D121) and S2 (D135, E137 and E139) motifs of full-length a-Syn (Fig. 2E). Furthermore, both S1 or S2 peptides reduce the CSPs of ^15^N-labeled a-Syn to a lesser extent than the addition of the a-Syn C-terminal peptide or the pS129 a-Syn C-terminal peptide. This suggests that S1 and S2 individually have a lower binding affinity for hTPR2A as compared to peptides that are comprised of both S1 and S2 (Fig. 2E). Taken together, these results show that both S1 and S2 compete for a single binding interface on hTPR2A.

**Figure 2.**
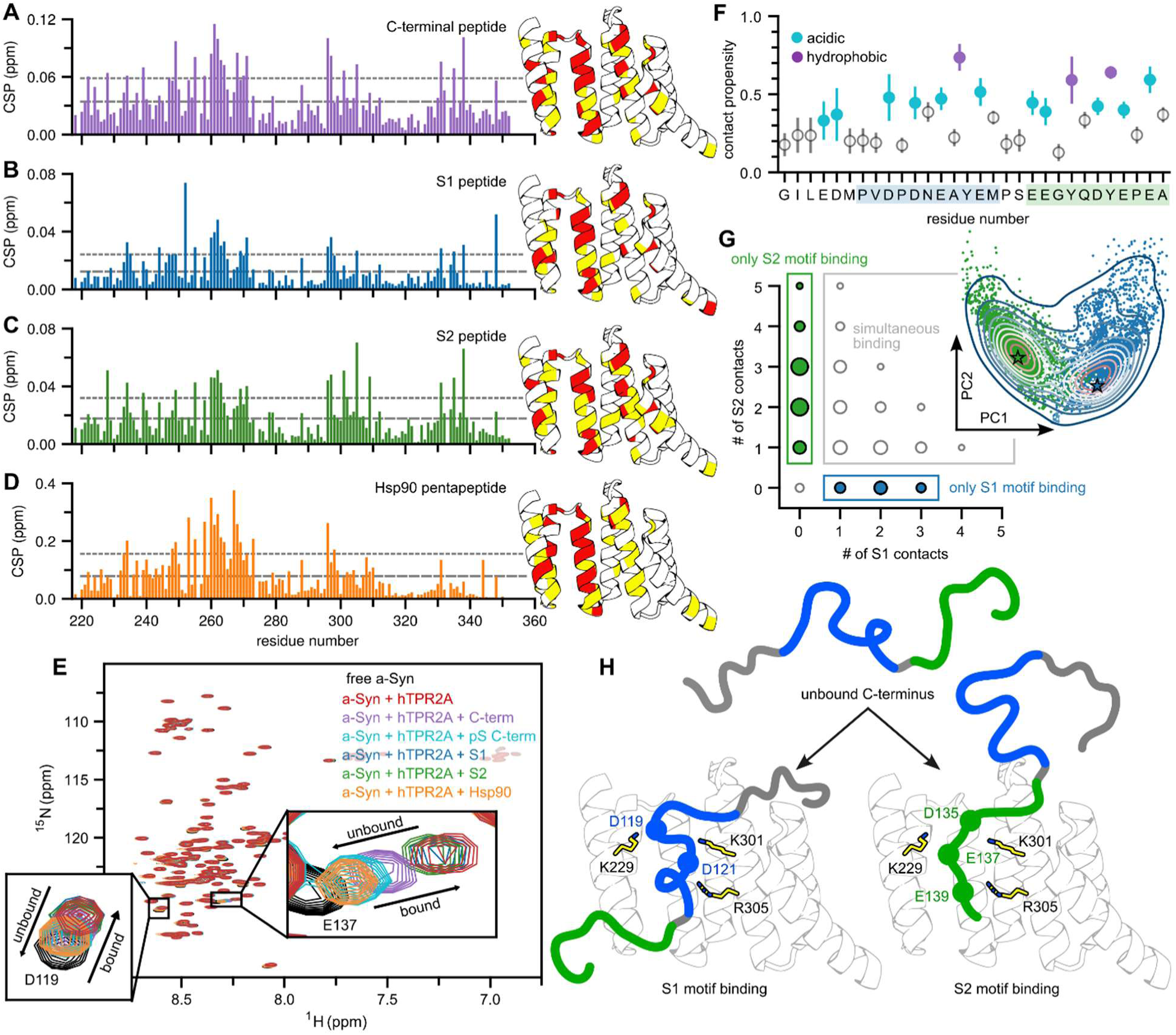
Competition for the Hsp90 pentapeptide interface on hTPR2A blocks binding of both S1 and S2 of a-Syn. (A-D) Quantification of chemical shift perturbations (CSPs) observed in residues of 200 µM ^15^N-labeled hTPR2A upon addition of 400 µM C-terminal a-Syn peptide (A), 800 µM S1 peptide (B), 400 µM S2 peptide (C) and 400 µM Hsp90 pentapeptide (D). Missing bars indicate prolines for which no chemical shift perturbations are measured. The dashed line represents the average CSP value, and the dotted line represents one standard deviation (SD) above the average CSP value. Residues which experience a CSP greater than the average are colored yellow and residues which experience a CSP greater than one SD above the average are colored red on the cartoon of the solved structure of hTPR2A bound to the Hsp90 pentapeptide (PDB:1ELR). (E) NMR competition assay monitoring the chemical shifts of 100 µM ^15^N-labeled a-Syn (black) in the presence of 100 µM hTPR2A (red) and the transitions between the bound and unbound states upon the addition of 100 µM a-Syn C-terminal peptide (purple), 100µM pS129 a-Syn C-terminal peptide (cyan), 100 µM S1 peptide (blue), 100 µM S2 peptide (green), or 100µM Hsp90 pentapeptide (orange). (F) Analysis of contact propensity for a-Syn residues with any other residues in hTPR2A. Acidic residues are colored cyan and aromatic side chain residues are colored purple. The S1 and S2 motifs are highlighted, in blue and green respectively. (G) The probability histogram of contact configurations between the S1 and S2 motifs and hTPR2A. Residues can form multiple contacts. Circle radius indicates the probability of finding a specific contact configuration e.g., top left marker corresponds to the probability of configurations with 0 S1 contacts and 5 S2 contacts. (G, inset) Electrostatic contact Principal Component Analysis (PCA) projection of all configurations (points) in the simulation ensemble. PC1 captures ∼40% of the variance. Points are continuously colored according to whether the configuration mainly involves interactions with S1 (blue), S2 (green), or both motifs (white). (H) Model of the interaction between S1 and S2 of a-Syn with hTPR2A highlighting key electrostatic interactions between S1 and S2 of a-Syn with the Hsp90 interacting residues (K229, K301, and R305) of hTPR2A.

### The S1 and S2 binding motifs of a-Syn compete for the Hsp90-binding pocket of hTPR2A

The TPR2A domain of STIP1/HOP mediates Hsp90 interactions by binding to its C-terminus [70]. Interestingly, our previous experiments demonstrated that the pentapeptide (MEEVD) comprising the C-terminus of Hsp90 is sufficient to disrupt mTPR2A/a-Syn binding [36]. We demonstrated here that many of the residues within hTPR2A that show CSPs upon binding a-Syn are adjacent to residues essential for its interaction with Hsp90. Upon addition of the Hsp90 pentapeptide, three clusters of residues in hTPR2A showed the largest CSPs, indicating interacting residues (Fig. 2D). Notably, the majority of the residues within these three clusters also show the largest CSPs upon addition of the C-terminal peptide of a-Syn and both the S1 and S2 peptides (Figs. 2A-C). Furthermore, competitive binding NMR experiments reveal that in the presence of equimolar hTPR2A, CSPs for residues in both S1 and S2 of ^15^N-labeled a-Syn are substantially reduced upon the addition of equimolar Hsp90 pentapeptide (Fig. 2E). Taken together, these results demonstrate that the Hsp90 pentapeptide interacts with the same interface on hTPR2A shared by both S1 and S2 of a-Syn and can displace S1 and S2 upon binding hTPR2A.

### Computational modelling suggests dynamic binding between a-Syn and hTPR2A

To model the multi-site interactions between a-Syn and hTPR2A, we performed extensive unbiased molecular dynamics (MD) simulations of the C-terminal peptide of a-Syn in the presence of hTPR2A. Global contact analysis suggested the importance of electrostatic interactions and interactions with residues containing aromatic side chains (Fig. 2F). To assess how S1 and S2 of a-Syn differentially interact with hTPR2A, we determined the time-dependent contact propensity of acidic residues in both motifs with positively charged residues in hTPR2A. Performing a principal component analysis (Fig. 2G, inset) we found ∼40% of the variance could be described by PC1 which also well separated the propensity for S1 (blue) or S2 (green) to form contacts. Quantifying the nature of these contacts revealed that although S1 and S2 can simultaneously bind to hTPR2A, nearly half of the contacts involved exclusively either S1 or S2, with S2 showing a higher propensity for these solo interactions (i.e., ∼26% vs. ∼16%) (Fig. 2G). Notably, the MD simulations predict that when S1 occupies the Hsp90 pentapeptide binding interface on hTPR2A, S2 can still make transient interactions with neighboring charged residues on hTPR2A and vice versa (Movie S1). The average number of total contacts for S1 was 2.34 ± 0.24, while for S2 was 2.63 ± 0.26. Overall, the simulations agree with our NMR findings, underlining the importance of electrostatic interactions in facilitating a-Syn/hTPR2A binding. The NMR analyses and MD simulations together suggest a dynamic binding mechanism between a-Syn and hTPR2A via the S1 and S2 motifs of a-Syn (Fig. 2H).

### Competitive two-site binding enhances the affinity of a-Syn for hTPR2A

The C-terminal a-Syn peptide was significantly more effective than the S1 and S2 peptides in competing with the full-length a-Syn for binding to hTPR2A. This suggests that multivalent interactions between the S1 and S2 motifs of a-Syn and hTPR2A increase the overall affinity of binding through avidity effects [71]. We employed Bio-Layer Interferometry (BLI) to determine the binding affinities of different a-Syn peptides for immobilized hTPR2A (Figure S4). The global binding affinity between hTPR2A and the C-terminal a-Syn peptide was determined to be K_d_ = 1.3 ± 0.4 µM (K_d_ = 7.9 ± 1.1 µM for immobilized C-terminal a-Syn peptide), which is significantly tighter than the binding affinity estimated by fitting the observed NMR CSPs with a 1:1 single-site binding model (K_d_ = 169 ± 67 µM), which does not account for the multivalent interaction involving the S1 and S2 motifs. We also measured the binding affinities of the S1, S2, and Hsp90 peptides for immobilized hTPR2A. In agreement with the results of the NMR competition experiments, the binding affinity of S2 (K_d_ = 32 ± 0.8 µM) is higher than that of S1 (K_d_ = 87 ± 7.8 µM), while the K_d_ of the Hsp90 pentapeptide for hTPR2A was 9.2 ± 0.7 µM (Figure S4), which agrees with a previous report (K_d_ ≈ 11 µM) [70]. Together, these results demonstrate that the dynamic interaction between S1 and S2 of a-Syn and hTPR2A significantly enhance the binding affinity in comparison to S1 and S2 separately.

### HOP inhibits a-Syn fibril formation

Our previous work has demonstrated that alteration of STIP1 expression in mouse models of α-synucleinopathy modulates the levels of insoluble pS129 a-Syn [36], suggesting that STIP1 may change the aggregation of a-Syn. Here, we further investigate if the direct interaction between STIP1 and a-Syn changes a-Syn aggregation *in vitro*. To this end, we conducted Thioflavin T (ThT) fluorescence assays to monitor a-Syn fibril formation. In the absence of STIP1, ThT fluorescence increases after an initial lag phase (Fig. 3A) as amyloid fibrils of a-Syn nucleate and begin to elongate as shown before [72–74]. Addition of STIP1 reduced ThT fluorescence in a concentration dependent manner, with sub-stoichiometric concentrations of STIP1 sufficient to significantly reduce ThT fluorescence (Fig. 3A). Notably, the addition of mTPR2A reduced the ThT fluorescence of a-Syn to the same extent as full-length STIP1 at equimolar concentrations, indicating that the mTPR2A domain alone is sufficient to attenuate a-Syn fibrilization (Fig. 3A). To exclude the possibility that the changes in ThT fluorescence are caused by non-specific crowding effects, we aggregated a-Syn in the presence of BSA at an equal mg/ml ratio as mTPR2A at a 1:1 a-Syn to mTPR2A concentration, which resulted in only minimal reduction of ThT fluorescence (Figure S5). Furthermore, we measured a-Syn fibril formation in the presence of the TPR1 domain of STIP1 (mTPR1, which has only minimal interactions with a-Syn [36]). NMR analysis of ^15^N-labeled a-Syn in the presence of mTPR1 shows only minor peak shifts, indicating that mTPR1 interacts with a-Syn at very low affinity, likely in a non-specific manner (Figure S6). The addition of mTPR1 reduced ThT fluorescence to a much lesser extent compared to mTPR2A at similar stoichiometric ratios, confirming that the effects observed for mTPR2A are not due to non-specific binding or crowding effects (Figs. 3B and 3C). Similarly, the addition of sub-stoichiometric concentrations of hTPR2A also reduced the final steady-state of a-Syn ThT fluorescence in a concentration dependent manner (Figs. 3D and 3E). Furthermore, both mTPR2A and hTPR2A substantially extended the lag phase of a-Syn fibril formation (Fig. 3F). In addition, when we separated a-Syn aggregates into soluble and insoluble fractions by centrifugation, the proportion of soluble a-Syn increases in the presence of hTPR2A (Figure S7). Taken together, these results suggest that the direct interaction between the TPR2A domain of STIP1/HOP and the C-terminal domain of a-Syn stabilizes a-Syn in conformations that favour more soluble, non-fibrillar species.

**Figure 3.**
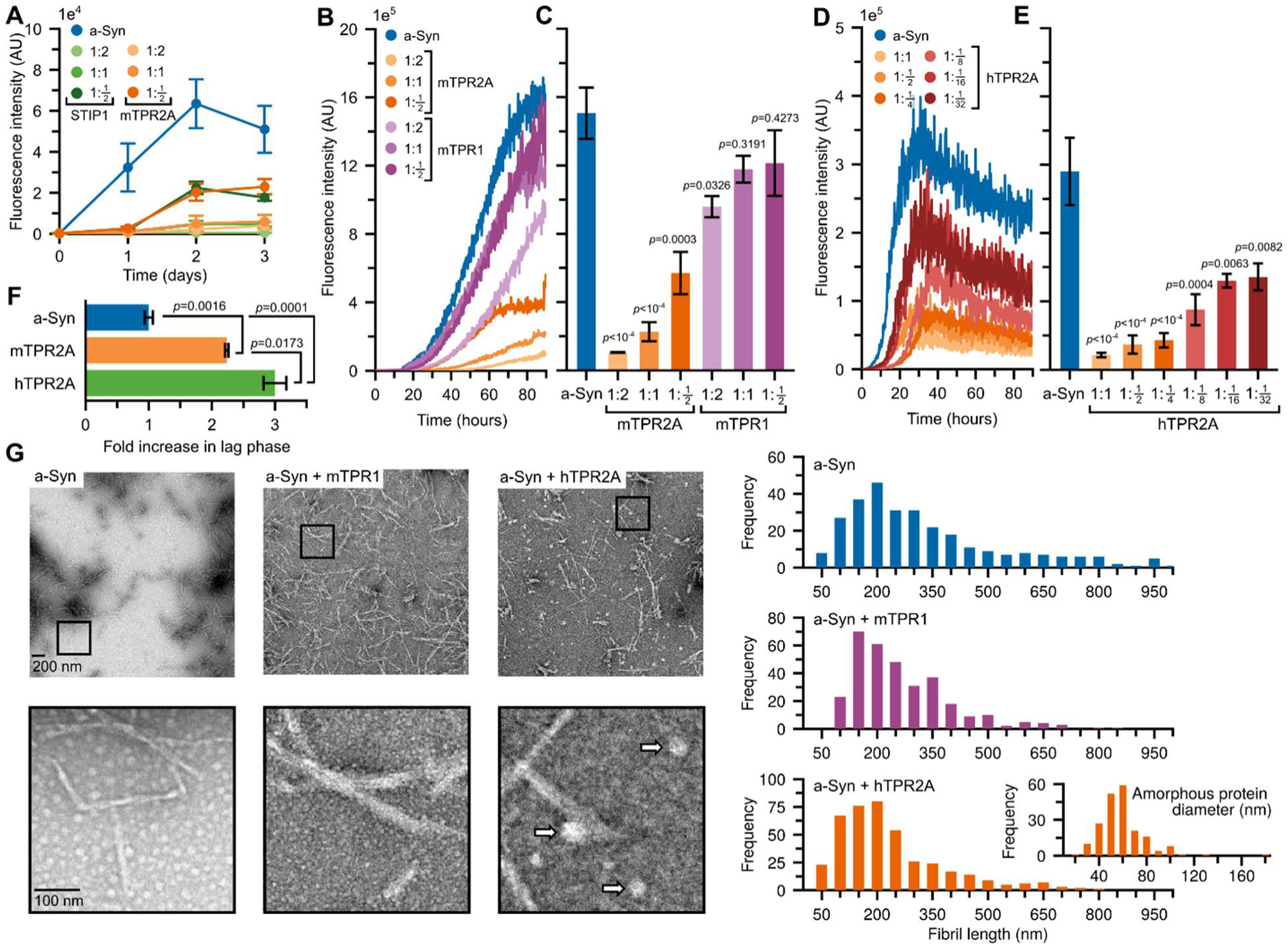
ThT analysis of a-Syn fibril formation in the presence of TPR2A. (A) Fibril formation of a-Syn (20 µM) measured by ThT fluorescence in the absence (blue) or presence of either STIP1 or the TPR2A domain of STIP1 at concentrations ranging from 1/2-fold to 2-fold relative to a-Syn. (B) Fibril formation of a-Syn (60 µM) measured by ThT fluorescence in the presence of mTPR2A or mTPR1 at concentrations ranging from 1/2-fold to 2-fold relative to a-Syn and (C) quantification of final steady-state values. (D) Fibril formation of a-Syn (60 µM) measured by ThT fluorescence in the presence of hTPR2A at concentrations ranging from 1/32-fold to 1-fold relative to a-Syn and (E) quantification of final steady-state values. (F) Comparison of lag phase extension when a-Syn (60 µM) is aggregated in the presence of equimolar mTPR2A or hTPR2A. Error bars represent standard errors of the mean. Statistical significance is in reference to the control condition unless otherwise indicated. (G) Representative EM images at low (upper) and high (lower) magnification taken after 5 days of a-Syn (175 µM) aggregation in the absence or presence of equimolar mTPR1 and hTPR2A with quantification of the distribution of fibril lengths observed and the diameter of the amorphous protein species observed in the presence of hTPR2A (highlighted by white arrows). To determine statistical significance, unpaired t-tests were used to compare means and standard deviations between relevant controls and experimental data sets (n=4). All data are mean ± SEM.

We next asked if hTPR2A binding to a single site in the C-terminus of a-Syn, i.e., either S1 or S2, is sufficient to alter fibril formation of a-Syn. We added hTPR2A to aggregation reactions containing a-Syn bearing alanine substitutions at D119 and D121 (ΔS1) or at D135, E137, and E139 (ΔS2). The addition of hTPR2A significantly reduced the ThT fluorescence of both ΔS1 or ΔS2 a-Syn (Figure S8), demonstrating that hTPR2A can attenuate a-Syn fibril formation by interacting independently with either the S1 or S2 motif within the C-terminus of a-Syn.

### hTPR2A promotes the accumulation of amorphous a-Syn aggregates

Since the overexpression of STIP1 in synucleinopathy mouse models increases the level of insoluble phosphorylated a-Syn [36], we speculated that a-Syn initially accumulates in ThT-negative forms in the presence of hTPR2A before eventually converting into fibrils. To test this, we analyzed a-Syn incubated with hTPR2A by Electron Microscopy (EM). Note that the samples contained a significantly higher concentration of a-Syn (175 µM) compared to our ThT experiments to allow efficient fibril visualization. The fibrils observed after a-Syn was aggregated alone for 5 days (Avg: 339 nm, Median: 272 nm) or in the presence of equimolar mTPR1 as a non-specific binding control (Avg: 269 nm, Median: 231 nm) are longer than those formed in the presence of hTPR2A (Avg: 246 nm, Median: 203 nm) (Fig. 3G). Furthermore, in the presence of hTPR2A, amorphous protein inclusions averaging ∼59 nm in diameter were observed, often in close proximity to fibrils (Fig. 3G, white arrows), which were not observed with a-Syn alone or treated with mTPR1. Notably, hTPR2A did not form these amorphous aggregates in the absence of a-Syn (Figure S9). Our observations suggest that the addition of hTPR2A attenuates fibril elongation by promoting either the formation or stabilization of amorphous high molecular weight a-Syn species.

### hTPR2A Modulates the Early Stages of a-Syn Aggregation

We next asked which stage of a-Syn fibril formation is affected by hTPR2A, specifically whether it inhibits nucleation, stalls elongation, or disassembles existing fibrils. We added equimolar hTPR2A to a-Syn fibrilization reactions prior to initiation of aggregation (0h), during late lag phase (6h), elongation phase (12h & 24h), and steady-state phase (48h). The addition of hTPR2A during late lag phase minimally extended the lag phase and reduced the ThT fluorescence during elongation, but did not alter steady-state ThT fluorescence after prolonged elongation (Figs. 4A-C). By contrast, hTPR2A added during the elongation phase or steady-state phase did not alter ThT fluorescence (Figs. 4A-C). By far the strongest reduction effects on a-Syn fibrilization occurred when hTPR2A was added prior to initiation, i.e., during the earliest phases of a-Syn aggregation.

**Figure 4.**
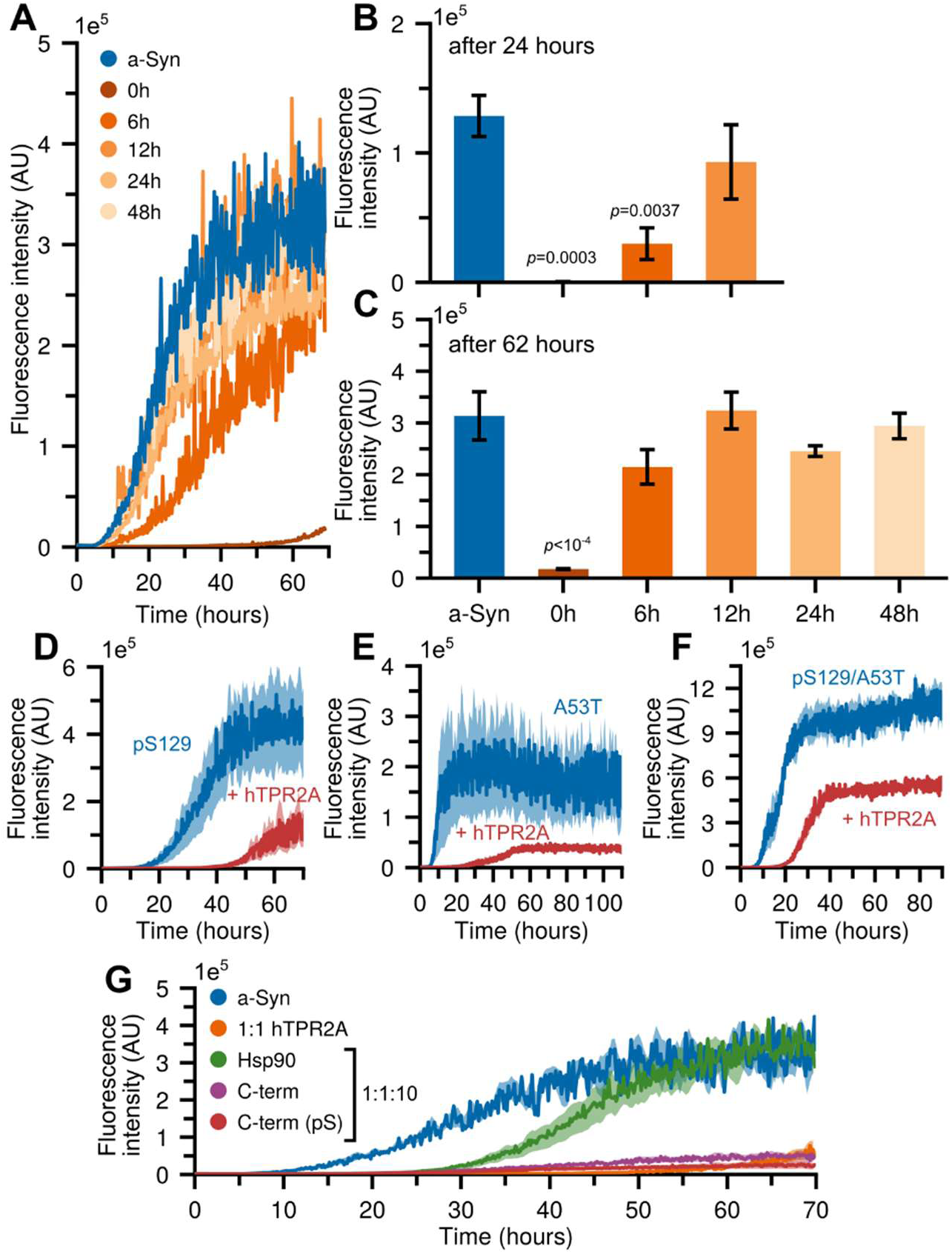
The addition of hTPR2A during early stages of a-Syn aggregation decreases the fibril formation of a-Syn and disease associated a-Syn variants. (A-C) Fibril formation of a-Syn (60 µM) measured by ThT fluorescence in the presence of hTPR2A at equimolar concentrations added at time 0, 6, 12, 24, and 48 hours after starting aggregation (A) and quantification of steady-state values at 24 hours (B) and 62 hours (C) of aggregation. Error bars represent standard errors of the mean. Statistical significance is in reference to the control condition unless otherwise indicated. (D-F) Fibril formation of pS129 a-Syn (D), A53T a-Syn (E), and pS129/A53T a-Syn (F) (60 µM) measured by ThT fluorescence in the presence of hTPR2A at equimolar concentrations. (G) Fibril formation of a-Syn (60 µM) measured by ThT fluorescence in the presence of equimolar hTPR2A and various peptides at a concentration of 10-fold excess of that of a-Syn. To determine statistical significance, unpaired t-tests were used to compare means and standard deviations between relevant controls and experimental data sets (n=4). All data are mean ± SEM.

To further test if hTPR2A inhibits a-Syn fibril elongation in seeded aggregation assays, we introduced additional a-Syn, either with or without hTPR2A, to aggregation reactions after the ThT fluorescence reached steady-state. The addition of a-Syn increased the ThT fluorescence beyond its previous steady-state value even in the presence of hTPR2A, indicating that hTPR2A did not prevent incorporation of a-Syn into existing fibrils (Figure S10), supporting the notion that hTPR2A influences a-Syn aggregation early, prior to fibril formation. Furthermore, hTPR2A is unable to disassemble fibrils, such as the disaggregase Hsp104 [75], as its addition at steady-state, i.e., after fibril formation, did not reduce ThT fluorescence.

Next, we investigated if addition of hTPR2A also inhibits fibril formation of a-Syn variants such as pS129 and A53T a-Syn. The addition of equimolar concentrations of hTPR2A to pS129 a-Syn aggregation reactions extended the lag phase and reduces the final steady-state value of ThT fluorescence (Fig. 4D). We obtained similar results with the PD-associated A53T mutant (Fig. 4E). Also, we found that the phosphorylation of A53T a-Syn at S129 did not prevent hTPR2A from reducing the fibrilization of a-Syn (Fig. 4F).

### Binding of the Hsp90 pentapeptide to hTPR2A inhibits its ability to modulate a-Syn aggregation

We next investigated whether blocking the interaction between hTPR2A and a-Syn abolishes the influence of hTPR2A on a-Syn aggregation. Since the Hsp90 pentapeptide (MEEVD) and the C-terminal a-Syn peptide can effectively compete with the full-length a-Syn for hTPR2A binding, we measured a-Syn fibril formation in the presence of equimolar hTPR2A and excess peptides (10-fold molar ratio). The addition of excess C-terminal or pS129 C-terminal a-Syn peptides to mixtures of a-Syn and hTPR2A shortened the lag phase of fibril formation by 45.9% (*p* = 0.0006) and 41.4% (*p* = 0.0012) respectively but did not prevent hTPR2A from reducing the steady-state ThT fluorescence of a-Syn (Fig. 4G). On the other hand, the addition of excess Hsp90 pentapeptide shortened the lag phase by 59.6% (*p* = 0.0004) and produced a similar maximum ThT fluorescence as a-Syn alone, suggesting that excess Hsp90 can mitigate hTPR2A regulation of a-Syn fibrilization (Fig. 4G). It is noteworthy that NMR ^1^H-^15^N HSQC analysis of ^15^N-labeled a-Syn in the presence of excess Hsp90 pentapeptide indicates that there are no direct interactions between the peptide and a-Syn (Figure S11). Taken together, these results suggest that blocking the binding of both S1 and S2 in the C-terminus of a-Syn to hTPR2A eradicates its inhibition of a-Syn aggregation.

### hTPR2A induces the formation of amorphous high molecular weight a-Syn species

Our results suggest that the interaction between hTPR2A and the C-terminal domain of a-Syn influences the pre-fibrillar stages of a-Syn aggregation. We next employed Atomic Force Microscopy (AFM) to further characterize the morphology of pre-fibrillar a-Syn species. We visualized samples taken at different time points from aggregation reactions containing 175 µM a-Syn in the absence or presence of equimolar mTPR1 or hTPR2A by AFM. The a-Syn fibrils formed in the presence of hTPR2A are shorter in length compared to reaction carried out in the presence of mTPR1 or a-Syn alone (Figure S12A). Moreover, fibril load (measured as the percentage of the surface area occupied by fibrils) of a-Syn samples aggregated in the presence of hTPR2A are significantly reduced compared to a-Syn aggregated alone or in the presence of mTPR1 (Figure S12B). Furthermore, we observed amorphous, high molecular-weight a-Syn species in samples aggregated in the presence of hTPR2A (Figure S12A). We performed aggregation reactions with reduced a-Syn concentrations to decrease the rate of fibril formation, allowing the capture and analysis of oligomeric species. The a-Syn oligomers visualized by AFM were transient, appeared after 2-3 days and disappeared upon fibril nucleation (day 3-4). However, in the presence of hTPR2A, many amorphous species were present after 1 day of aggregation and can be observed at all subsequent time points (Fig. 5A). The amorphous a-Syn species formed in the presence of hTPR2A are of a broad size range and are significantly larger than the oligomers measured in a-Syn samples aggregated alone or in the presence of mTPR1, as observed on the 3^rd^ day of aggregation (Figs. 5B and 5C). While oligomers of a-Syn were small and transient, the amorphous a-Syn species formed in the presence of hTPR2A persisted for several days and increased in size, as is apparent by correlation plots and distributions of their diameter and height (Figure S12C).

**Figure 5.**
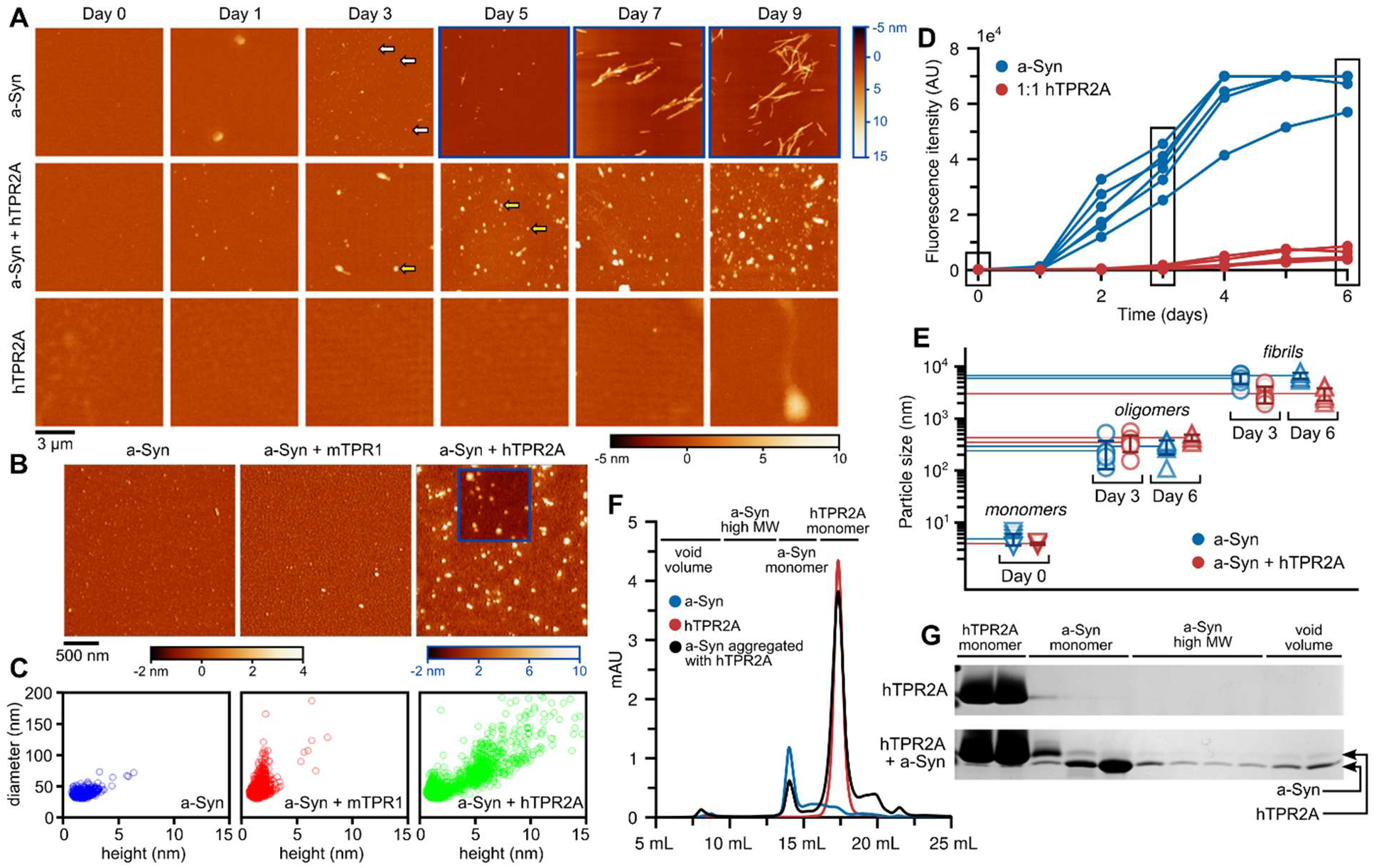
hTPR2A promotes the formation of and maintains large amorphous a-Syn species. (A) Representative AFM images of a-Syn aggregation (60 µM) in the absence and presence of and hTPR2A at an equimolar concentration as a function of time. Protein samples are deposited on a mica surface. Scale bar applies to all images. The color map with black font corresponds to all images except those out lined in blue. The color map with blue font corresponds to those images. Grey arrows indicate a-Syn oligomers and yellow arrows indicate what we define here as amorphous high molecular weight aggregates. (B) Representative AFM images of amorphous high molecular weight a-Syn species produced during the aggregation of a-Syn in the absence or presence of mTPR1 and hTPR2A at equimolar concentrations taken on the 3^rd^ day of incubation when the population of a-Syn oligomers was largest. (C) Correlation plots of the distribution of the heights and diameters of the amorphous high molecular weight a-Syn species produced in the absence or presence of mTPR1 and hTPR2A on the 3^rd^ day of incubation. (D) Fibril formation of a-Syn (20 µM) measured by ThT fluorescence in the absence (blue) or presence of equimolar (red) concentration of hTPR2A. Samples were collected after 3 and 6 days of aggregation for further DLS analysis. (E) Quantification of the hydrodynamic radius measured in nm of a-Syn monomers (4 nm), oligomers (100-800 nm), and fibrils (>1,000 nm) after 60 µM a-Syn was aggregated for 3 and 6 days at 37 °C in the absence (blue) or presence of hTPR2A (red), as determined using a DynaPro NanoStar dynamic light scattering instrument (Wyatt Technologies). (F) Chromatographs produced by passing a-Syn (300 µM) (blue), a-Syn with equimolar hTPR2A (black), or hTPR2A alone (red) that was incubated at 37 °C for 48 hours through a Superdex Increase 200 size exclusion column. Peaks corresponding to the location of monomeric hTPR2A, monomeric a-Syn, and high molecular weight oligomeric a-Syn species bound to hTPR2A are labeled on the chromatograph. (G) SDS-PAGE gel of fractions collected from the Superdex Increase 200 size exclusion column demonstrating the change in the fractionation of hTPR2A.

To verify the aggregate size differences, we monitored a-Syn species present in aggregated samples by Dynamic Light Scattering (DLS). Aligning with EM and AFM results, DLS analysis showed a significant reduction in the size of the fibrils formed in the presence of hTPR2A. Also, in agreement with AFM, the size of the oligomers formed in the absence of hTPR2A are significantly smaller than the pre-fibrillar a-Syn species formed in the presence of hTPR2A (Figs. 5D and 5E). Taken together, our results suggest that, in contrast to the oligomers formed by a-Syn alone, the interaction between hTPR2A and the C-terminal domain of a-Syn promotes the formation of larger amorphous a-Syn species that do not efficiently transition into fibrils. These accumulating amorphous a-Syn species formed in the presence of hTPR2A are not static and continue to increase in size as aggregation progresses (Fig. 5A and Figure S12A), suggesting that hTPR2A may stabilize soluble a-Syn aggregates thereby facilitating their continued growth rather than transition to fibrils.

### hTPR2A associates with high molecular weight species of a-Syn during aggregation

Molecular chaperones that interact with the N-terminal domain of a-Syn, such as Hsp90 and Hsp70, maintain their N-terminal interactions with oligomeric a-Syn [26]. To test if hTPR2A remains bound to the C-terminal domain of a-Syn within the amorphous high-molecular weight a-Syn species, we analyzed aggregated samples using size-exclusion chromatography. Samples of hTPR2A that were incubated at 37 °C for 24 hours in a shaking incubator before injecting onto a Superdex 200 10/300 size exclusion column elute as a sharp peak corresponding to monomeric hTPR2A (Fig. 5F). On the other hand, when hTPR2A is incubated with equimolar a-Syn, hTPR2A elutes in fractions corresponding to monomeric hTPR2A and co-elutes with a-Syn in fractions corresponding to monomeric and high molecular weight species of a-Syn (Figs. 5F and 5G). The co-elution of hTPR2A and a-Syn in high molecular weight fractions suggests that hTPR2A interacts with both monomeric a-Syn and a-Syn within the amorphous high molecular weight species.

### Co-expression of HOP and a-Syn in yeast alters a-Syn localization and increases a-Syn toxicity

To test if the interaction between a-Syn and HOP modifies a-Syn aggregation and toxicity within a cellular context, we employed a well-established yeast model of a-Syn toxicity [76,77]. Low-copy expression of a-Syn in yeast does not impart a growth defect at optimal growth conditions (30 °C) but reduces growth under heat stress conditions (37 °C) [76,78]. Co-expression of HOP and a-Syn unmasks a growth defect in yeast even in the absence of heat stress (Fig. 6A) and exacerbates the growth defect under heat stress conditions (Fig. 6B). Treatment with Hsp90 ATPase inhibitor Radicicol did not recover the growth phenotype observed with co-expression of HOP and a-Syn at either optimal (Fig. 6C) or heat stress temperatures (Fig. 6D), suggesting that the growth defect is due to the interaction between HOP and a-Syn, independent of Hsp90 ATP-hydrolysis.

**Figure 6.**
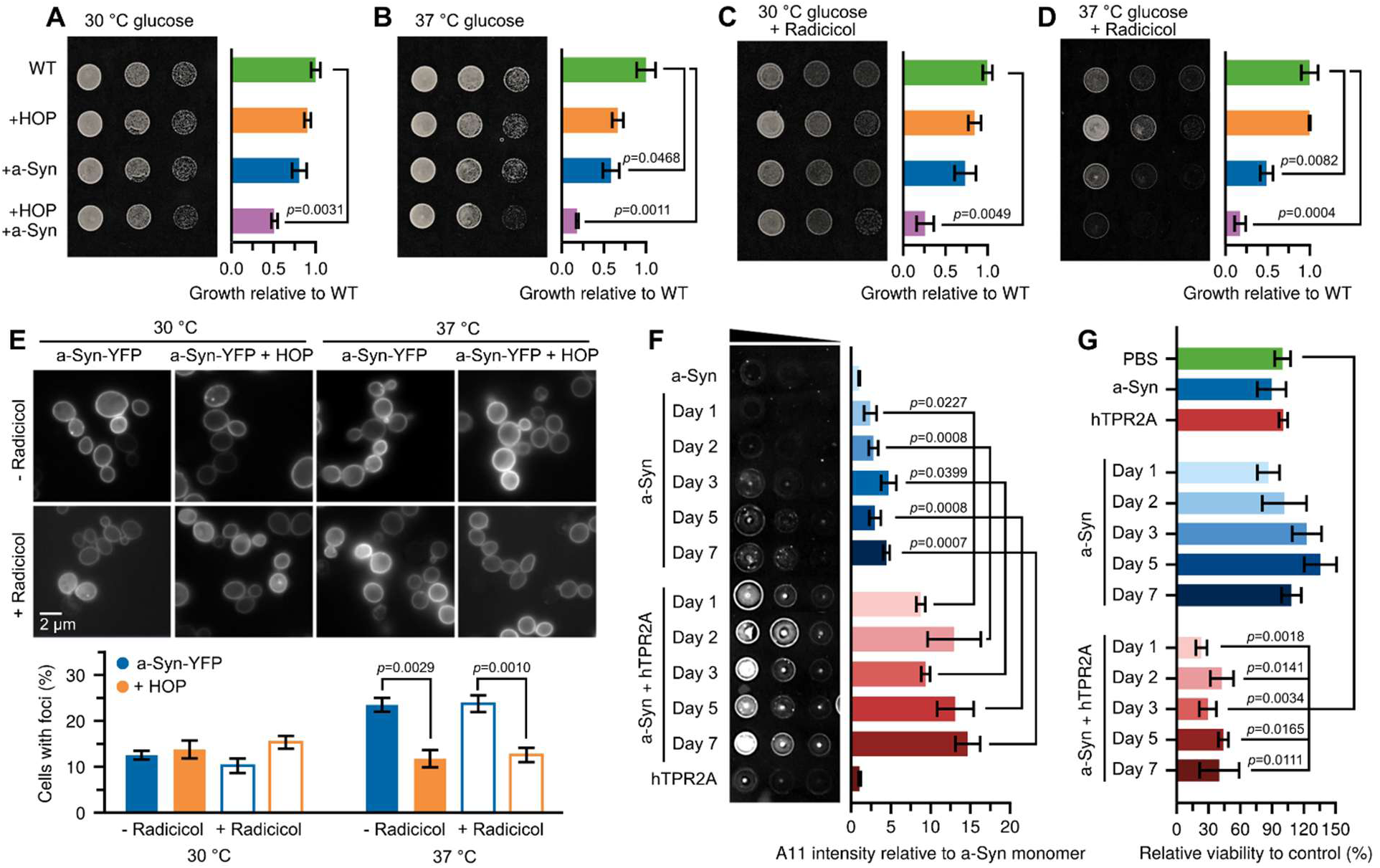
hTPR2A promotes amorphous oligomer-like a-Syn species and increases the toxicity of a-Syn. (A-D) Spotting assay of wildtype (BY4741) yeast expressing a control plasmid, HOP, a-Syn, or expressing HOP and a-Syn grown at optimal growth temperatures (30°C) (A, C) or under heat stress conditions (37 °C) (B, D) on SD media in the absence (A, B) or presence of Radicicol (C, D). (E) Upper: Fluorescence microscopy of yeast (BY4741) cells expressing a-Syn-YFP or co-expressing both a-Syn-YFP and HOP grown under optimal and heat stress growth temperatures in the absence and presence of Radicicol. Lower: Quantification of a-Syn cytoplasmic foci observed under each condition. (F) Samples of a-Syn aggregated in the absence or presence of equimolar hTPR2A from subsequent timepoints were spotted in three five-fold dilutions from left to right and examined by dot blotting analysis using the anti-oligomer antibody A11. The quantitative analysis of the A11 signal relative to monomeric a-Syn was performed for 3 independent aggregation reactions of each mixture. (G) Quantification of cell viability measured from SH-SY5Y cells after treatment with 35 µM of a-Syn aggregated in the absence or presence of equimolar hTPR2A for subsequent time periods and control samples of monomeric a-Syn, hTPR2A and PBS buffer. To determine statistical significance, unpaired t-tests were used to compare means and standard deviations between relevant controls and experimental data sets (n=3). All data are mean ± SEM.

To test if expression of HOP alters the localization of a-Syn *in vivo*, we monitored yeast cells expressing HOP and a-Syn-YFP from low copy number plasmids by fluorescent microscopy. In agreement with previous studies [76], we observed a-Syn-YFP localization at the plasma membrane and in concentrated cytoplasmic inclusions, which increase in number during heat stress (37 °C). Co-expressing HOP with a-Syn-YFP did not change the proportion of cells containing cytoplasmic a-Syn-YFP inclusions when grown under optimal conditions. However, it significantly reduced this proportion under heat stress conditions (Fig. 6E). To test whether the change in a-Syn localization is caused by the interaction between HOP and a-Syn or a passing of a-Syn to Hsp90, cells were treated with Radicicol at concentrations below the threshold to induce the heat shock response [79]. Treatment with Radicicol did not alter the effect of HOP co-expression in reducing the proportion of cells with cytoplasmic inclusions after heat stress (Fig. 6E), suggesting that HOP changes a-Syn-YFP localization independently of Hsp90 ATPase activity.

### Amorphous high molecular weight a-Syn species formed in the presence of hTPR2A contain cytotoxic a-Syn oligomers

Since HOP altered the toxicity of a-Syn when co-expressed in the yeast model system, we tested if the amorphous high molecular weight a-Syn species formed in the presence of hTPR2A resemble cytotoxic a-Syn observed in synucleinopathy models [80,81]. The oligomer content of samples from aggregation mixtures containing a-Syn with or without the addition of hTPR2A were quantified using immunodetection by the A11 antibody, which specifically recognizes pre-fibrillar a-Syn assemblies, including pathogenic a-Syn oligomers [80,81]. We applied a-Syn species formed at different time points upon incubation with hTPR2A to a membrane and probed with the A11 antibody. Significantly more A11-positive signal was detected in samples of a-Syn aggregated in the presence of hTPR2A across all time points measured as compared to reaction performed in its absence (Fig. 6F). These results suggest that the amorphous high molecular weight a-Syn species that accumulate in the presence of hTPR2A are comprised of or resemble the oligomeric a-Syn species that have been observed in synucleinopathy models [81,82]. Furthermore, the increased A11 signal confirms that a-Syn aggregated in the presence of hTPR2A forms increased levels of pre-fibrillar a-Syn species.

### hTPR2A exacerbates the toxicity of aggregated a-Syn on SH-SY5Y cells

Since smaller pre-fibrillar species are more toxic to neuronal cells than larger fibrillar a-Syn [83], we tested if a-Syn samples aggregated in the presence of hTPR2A are cytotoxic to cells by assessing the viability of SH-SY5Y human neuroblastoma cells treated with different aggregated samples of a-Syn. When cells were treated with samples from aggregation mixtures containing only a-Syn (35 µM initial monomer concentration), no reduction on cell viability was observed for any of the aggregation times tested. By contrast, treatment with samples from aggregation mixtures containing a-Syn and equimolar hTPR2A from all timepoints tested caused a significant reduction in cell viability as compared to cells topically treated with aggregated a-Syn alone (Fig. 6G). These results show that the accumulation of amorphous high molecular weight a-Syn species caused by hTPR2A binding increases sample toxicity relative to mostly fibrillar a-Syn aggregates.

## Discussion

Our study deciphers how the molecular co-chaperone STIP1/HOP prevents the fibrilization of a-Syn by steering it toward the formation of highly toxic oligomeric species. The two independent low-affinity binding motifs in the C-terminus of a-Syn compete for a single interface on the TPR2A domain of STIP1/HOP. This dynamic binding facilitates transient interactions between these two proteins and dictates the specificity for the TPR2A over the other TPR domains of STIP1/HOP.

Molecular chaperones and co-chaperones help maintain protein functions during misfolding stress [84]. Of note, however, they can also stabilize or perpetuate toxic conformations of misfolded proteins [24,25,85–87]. Our previous studies in mouse models found that STIP1 interacts with a-Syn and that decreased levels of STIP1 mitigate a-Syn toxicity and reduce the levels of S129 phosphorylated a-Syn, a neuropathological marker of synucleinopathies. Conversely, increased STIP1 levels exacerbate synucleinopathy-associated phenotypes and the accumulation of S129 phosphorylated a-syn [36]. Our biochemical analyses presented here reveal how the interaction between a-Syn and STIP1/HOP promotes the formation of oligomeric, non-fibrillar conformers of a-Syn. In addition, a-Syn retains its binding to STIP1/HOP within these amorphous aggregates, further attenuating the transition into fibrils and promoting the accumulation of amorphous oligomeric species. We have found previously that the binding of STIP1 enhances the S129 phosphorylation of a-Syn. The retained binding of STIP1/HOP to the amorphous aggregates of a-Syn may explain the accumulation of phosphorylated S129 a-Syn aggregates in the synucleinopathy mouse model with increased STIP1 levels [36].

We further show that STIP1/HOP does not require ATP-dependent chaperoning functions or other contributions by Hsp90 and Hsp70 to promote the accumulation of oligomeric a-Syn. Rather, STIP1/HOP modulates a-Syn aggregation through a holdase mechanism that only requires a direct interaction [88,89]. There are specific scenarios that may favour a holdase function of STIP1/HOP, such as in aged cells that are challenged by an ongoing proteostasis crisis [90,91], where STIP1/HOP favours client proteins over Hsp90 and Hsp70 [92]. This change in interaction preference may be a consequence of Hsp90 and Hsp70 being tied up in interactions with misfolded proteins [44,92] and ATP depletion, thereby favouring the holdase functions of STIP1/HOP [93–95]. In addition, both STIP1/HOP and a-Syn are found in extracellular vesicles [47,96,97] which can explain why a-Syn phosphorylated at S129 is enriched inside these vesicles [98]. When secreted into the ATP-deprived extracellular space, the holdase function of STIP1/HOP may alter the interaction between misfolded a-Syn and extracellular proteins, as observed between STIP1/HOP and PrP^C^ [99,100], thereby contributing to cell-to-cell spreading of misfolded a-Syn. Moreover, a-Syn oligomers have been previously shown to coopt PrP^C^, mGluR5 and NMDA receptors to cause synaptic toxicity, suggesting another mechanism by which interaction of STIP1 and a-Syn may act [35,101]. The holdase activity of STIP1/HOP generally mitigates the toxicity associated with misfolded proteins during proteostasis stress [43,44,102]. Yet, we demonstrate here for a-Syn that it can also have detrimental effects on aggregation-prone proteins.

Many molecular chaperones, such as Hsp90 and Hsp70, reduce the toxicity of a-Syn oligomers by binding a-Syn and masking regions of its N-terminal domain, thereby preventing membrane disruption and cellular toxicity [6,26–28,30]. Intriguingly, our data suggest that Hsp90 and a-Syn can compete for binding to STIP1/HOP, suggesting that the relative abundance and localization of each of these molecular chaperones may determine a-Syn toxicity. Our previous studies show that during protein misfolding stress, STIP1/HOP accumulates into defined cytoplasmic areas along with soluble ubiquitinated proteins [44]. These areas may favour the deleterious interaction between a-Syn and STIP1/HOP over the protective binding between a-Syn and Hsp90.

The same key acidic residues in the C-terminal domain of a-Syn which interact with STIP1/HOP also mediate binding to FABP3 [33], tau [103], and Ca^2+^ [104], all of which also promote a-Syn oligomerization [31–33,36]. These acidic residues are essential for recruiting monomeric and oligomeric a-Syn into existing fibrils [18], and alterations in the C-terminal charge of a-Syn modulate the kinetics of fibril formation [7–10]. Specific residues in the N-terminal domain of a-syn and the hydrophobic NAC region are key contact sites within a-Syn fibrils [105–107], whereas the negatively charged C-terminal domain remains disordered, forming a fuzzy coat around fibrils that facilitate the “seeding” process by recruiting a-Syn monomers and oligomers through their positively charged N-terminus [18].

With all these features in mind, we propose a general mechanism by which proteins interacting with the C-terminus of a-Syn masks its negative charge, thereby reducing the intra- and inter-molecular N-C terminal interactions. Consequently, binding of STIP1/HOP, FABP3, tau, and calcium ions to a-Syn may lower the kinetic barrier to form highly toxic oligomers and simultaneously prevent the conversion of oligomers into fibrils (Fig. 7). Thus, N-terminal interactions, such as those observed for Hsp90 and Hsp70, may prevent the accumulation of cytotoxicity a-Syn oligomers, whereas C-terminal interactions, like STIP1/HOP, promote the accumulation of cytotoxic a-Syn (Fig. 7). This baffling dichotomy deserves consideration in studies seeking to determine the effect of molecular chaperones and other proteostatic pathways on the toxicity of a-Syn in PD and LBD. Moreover, our findings may apply to other aggregation-prone proteins associated with neurodegenerative diseases and indicate a possible protective role of fibrillar, amyloid-like aggregates and detrimental effects of molecular chaperones that prevent fibrillar aggregation and favour the accumulation of highly toxic oligomers.

**Figure 7.**
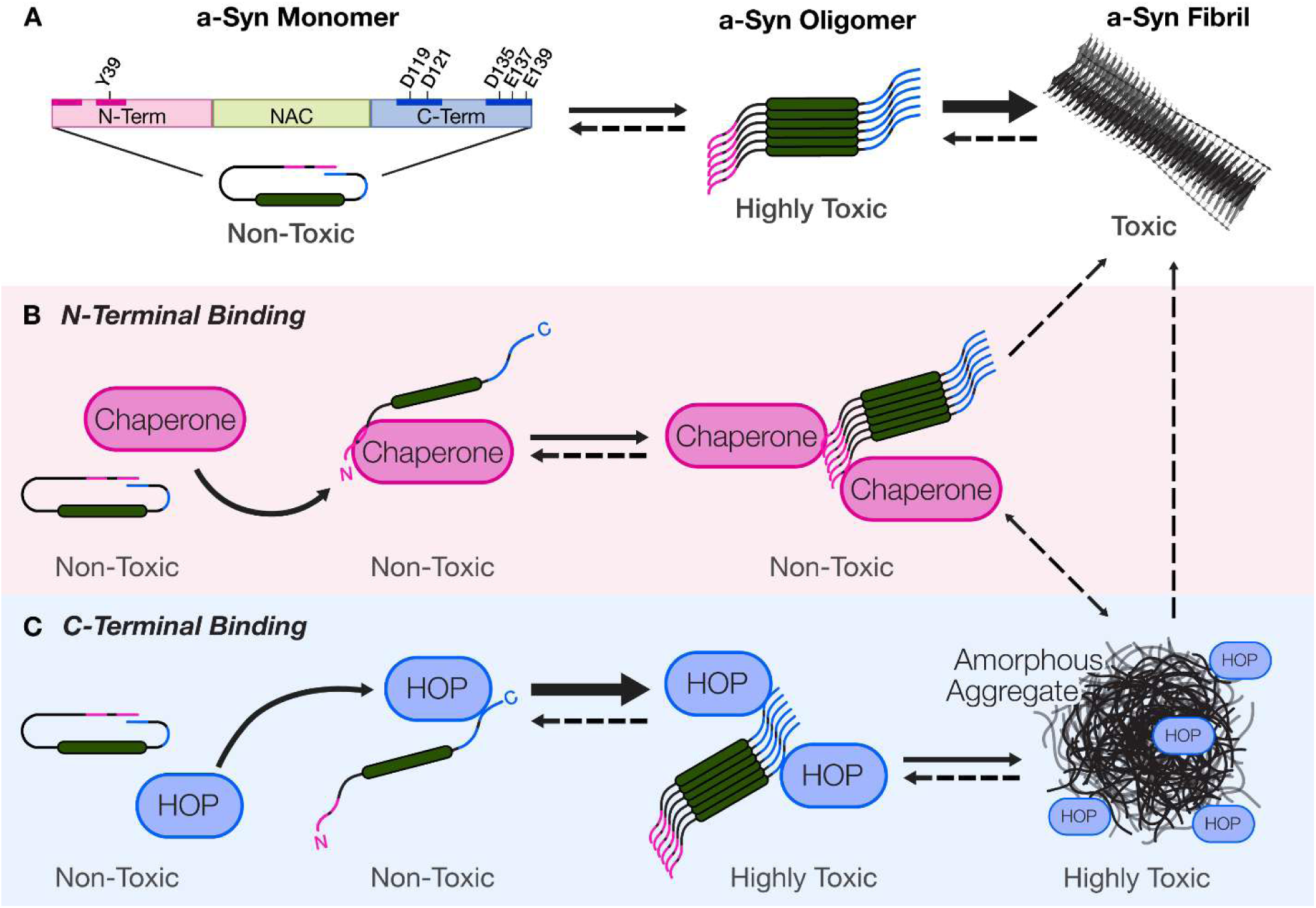
Protein interactions with the N- and C-terminal domains of a-Syn modulate aggregation and toxicity. (A) Intramolecular interactions between the terminal domains of monomeric a-Syn provide an energetic barrier to oligomer formation and favor fibril formation. (B) Molecular chaperones recognize and bind hydrophobic motifs (pink) in the N-terminal domain, maintain monomeric a-Syn conformers and masking the toxicity of oligomers. (C) HOP interacts with motifs of charged residues (blue) in the C-terminal domain of a-syn, promoting the formation of toxic amorphous aggregates.

## Data, Materials, and Software Availability

All study data are included in the article and/or Supplemental Information.

## Acknowledgments

We thank the Biomolecular NMR and BioCORE facilities (Western University) for their assistance and use of equipment. Digital Research Alliance of Canada provided the computational resources. Thank you to Karling Frankel for creating the digital design of our aggregation model schematic.

## Author Contributions

B.S.R., designed and performed research, analyzed data, generated figures, and wrote the paper. C.J.W. and J.L performed research, analyzed data, and aided in figure generation. R.M.L., J.C.J.C., and E.D.C.P. performed research and analyzed data. M.K., T.M.D., and E.A.F designed research and contributed analytical tools. M.A.M.P, J.L, M.L.D, and W.Y.C designed research, contributed reagents or analytical tools, and wrote the paper. All authors aided in revising the manuscript’s intellectual content. All authors read and approved the submitted manuscript.

## Ethics approval and consent to participate

N/A

## Consent for publication

N/A

## Abbreviations

a-Syn: alpha-synuclein
LBD: Lewy Body Dementia
PD: Parkinson’s disease
NAC: non-Aβ component
IDP: intrinsically disordered protein
FABP3: fatty acid binding protein 3
PrP^C^: cellular prion protein
STIP1: Stress inducible phosphoprotein 1
HOP: Hsp-organizing protein
TPR: tetratricopeptide repeat domain
DP: aspartate and proline rich domain
TDP-43: TAR DNA-binding protein 43
pS129 a-Syn: alpha-synuclein phosphorylated at serine 129
mTPR2A: mouse STIP1-TPR2A domain
hTPR2A: human HOP-TPR2A domain
NMR: Nuclear Magnetic Resonance
HSQC: Heteronuclear Single Quantum Coherence
CSPs: chemical shift perturbations
S1: residues 117-127 of alpha-synuclein
S2: residues 130-140 of alpha-synuclein
MD: molecular dynamics
BLI: Bio-Layer Interferometry
ThT: Thioflavin T
ΔS1: alpha-synuclein bearing alanine substitutions at D119 and D121
ΔS2: alpha-synuclein bearing alanine substitutions at D135, E137, and E139
EM: Electron Microscopy
AFM: Atomic Force Microscopy
DLS: Dynamic Light Scattering

