## Supplementary Information for "STIP1/HOP Promotes the Formation of Cytotoxic α-Synuclein Oligomers"

### Supplementary Figures and Video

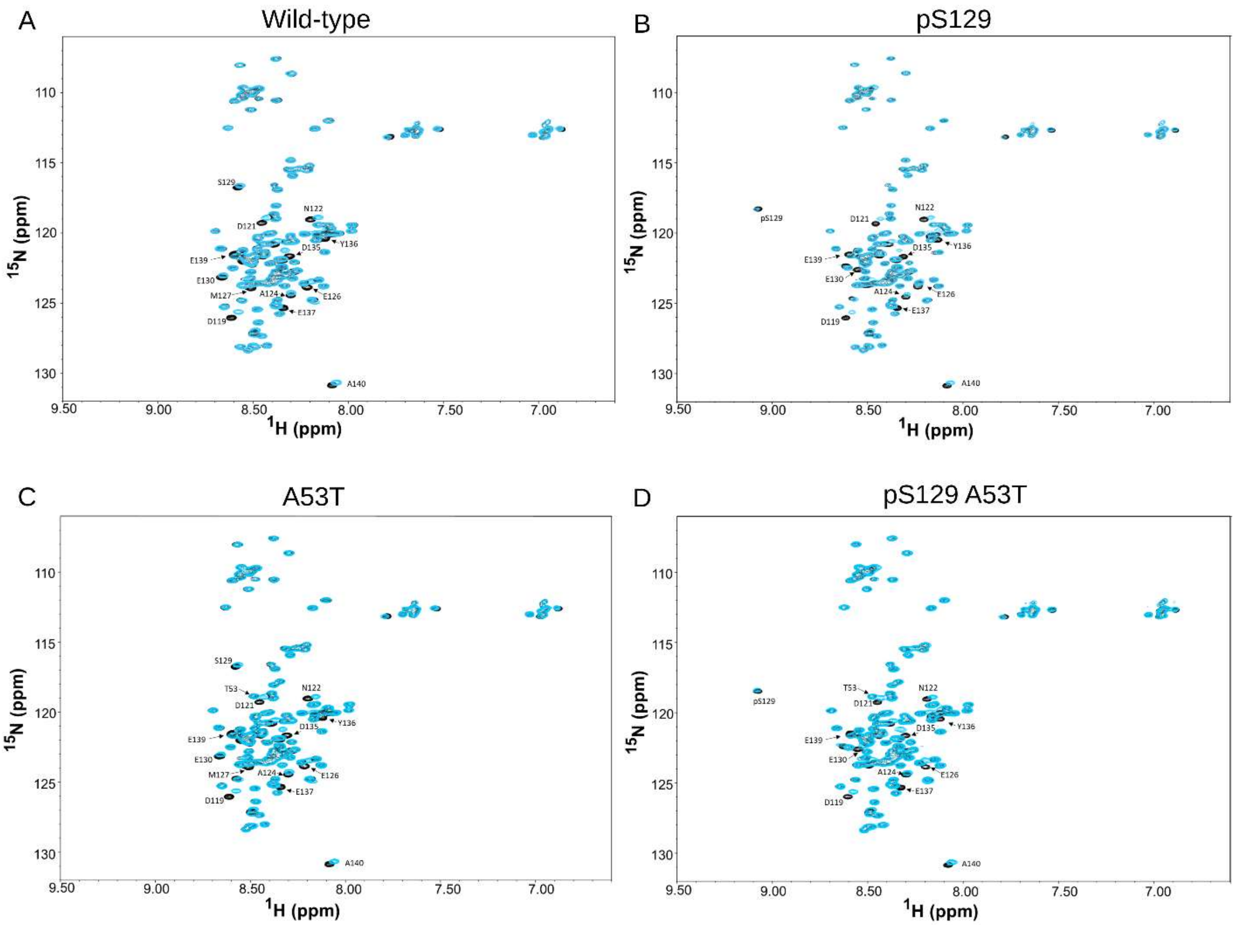


**Figure S1. The TPR2A domain of STIP1 mediates binding of a-Syn and its pathological variants.** $^{1}H$-$^{15}N$ HSQC spectra of 100 μM a-Syn (A), pS129 a-Syn (B), A53T a-Syn (C) and pS129 A53T a-Syn (D) in the absence (black) and the presence (cyan) of 200 μM of mTPR2A.

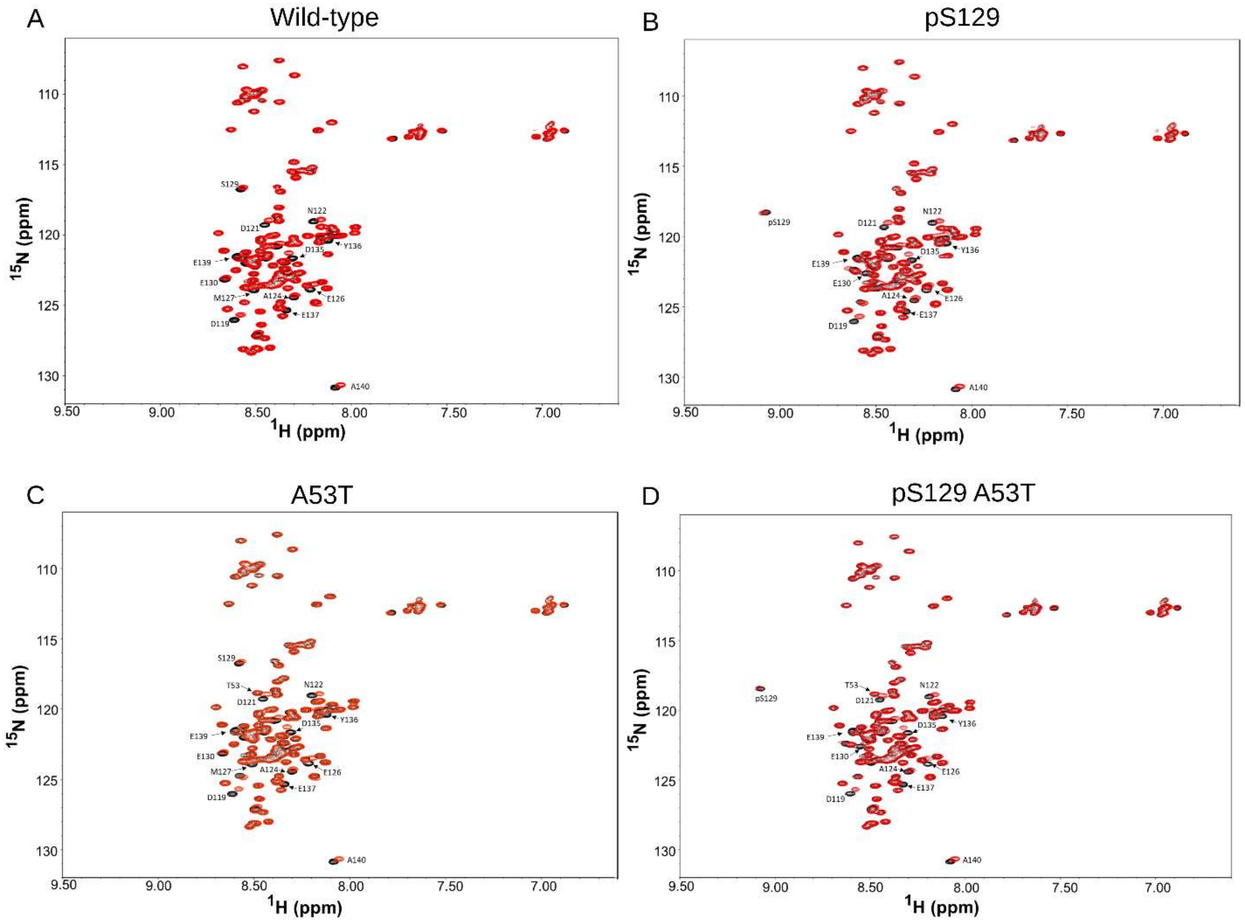


**Figure S2. The TPR2A domain of HOP mediates binding of a-Syn and its pathological variants.** $^{1}H$-$^{15}N$ HSQC spectra of 100 µM a-Syn (A), pS129 a-Syn (B), A53T a-Syn (C) and pS129 A53T a-Syn (D) in the absence (black) and the presence (red) of 200 µM of hTPR2A.

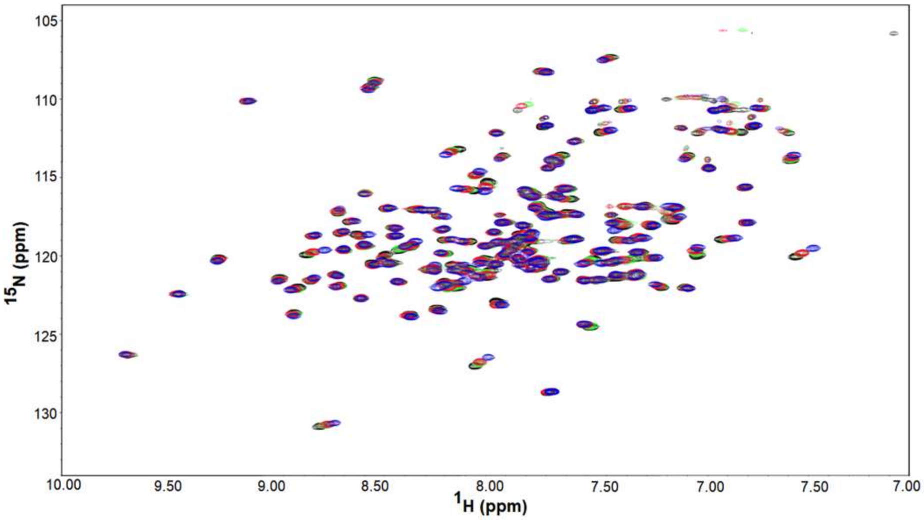


**Figure S3. Interacting residues of hTPR2A upon the addition of a-Syn peptides.** $^{1}$H-$^{15}$N HSQC spectra of 200 μM $^{15}$N-labeled hTPR2A (black) in the presence of 400 μM C-terminal a-Syn peptide (blue), 800 μM S1 peptide (green), and 400 μM S2 peptide (red).

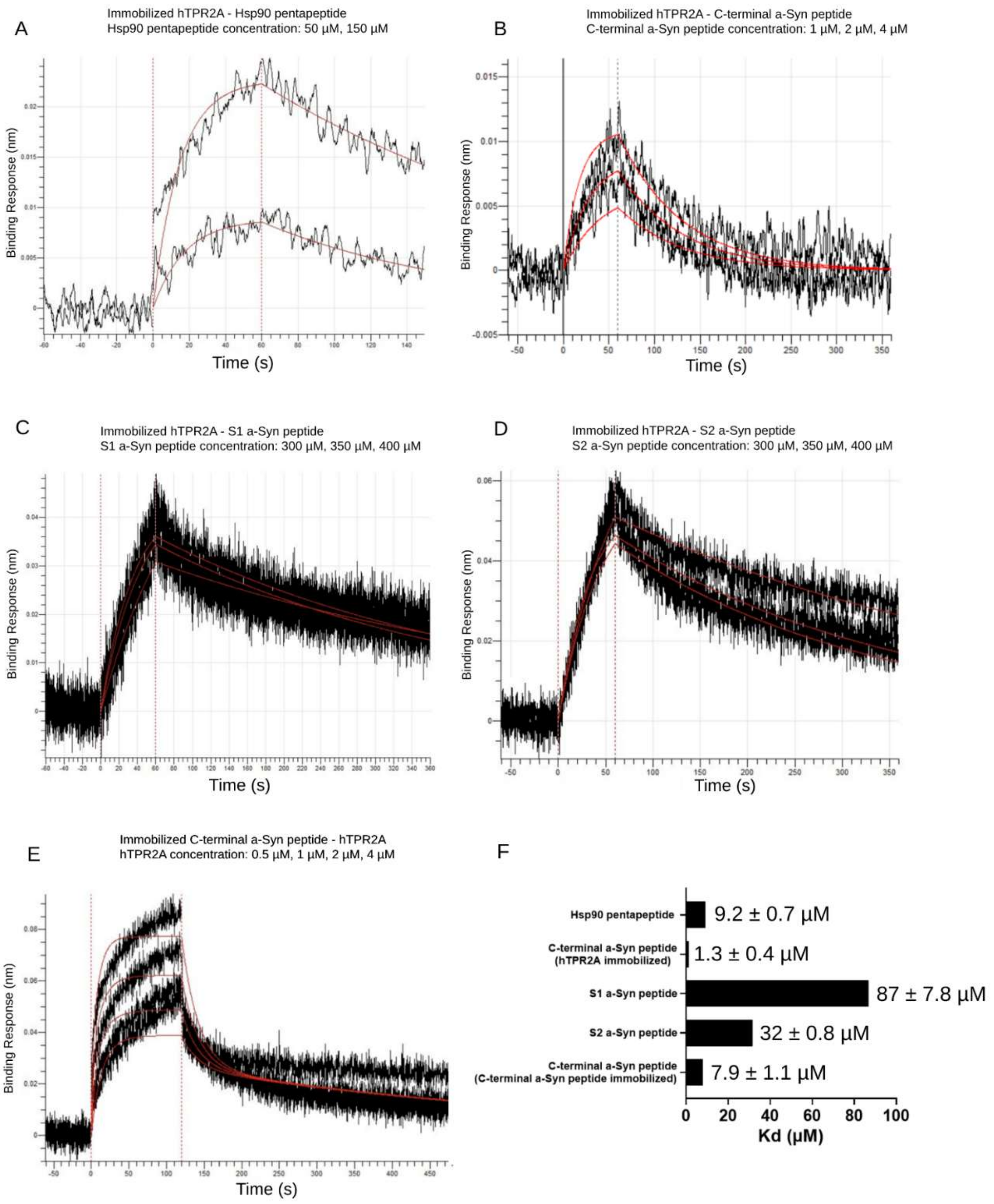


**Figure S4. The C-terminal a-Syn peptide binds hTPR2A with a higher affinity than S1 and S2 peptides.** BLI analysis of the binding of Hsp90 pentapeptide to immobilized hTPR2A (A), C-terminal peptide to immobilized hTPR2A (B), S1 a-Syn peptide to immobilized hTPR2A (C), S2 a-Syn peptide to immobilized hTPR2A (D), and hTPR2A to immobilized C-terminal peptide (E). Analytes were used at the indicated concentrations. Measured binding curves are depicted in black and fitted $K_d$ curves are depicted in red. (F) Quantification of $K_d$ values determined by kinetic analysis.

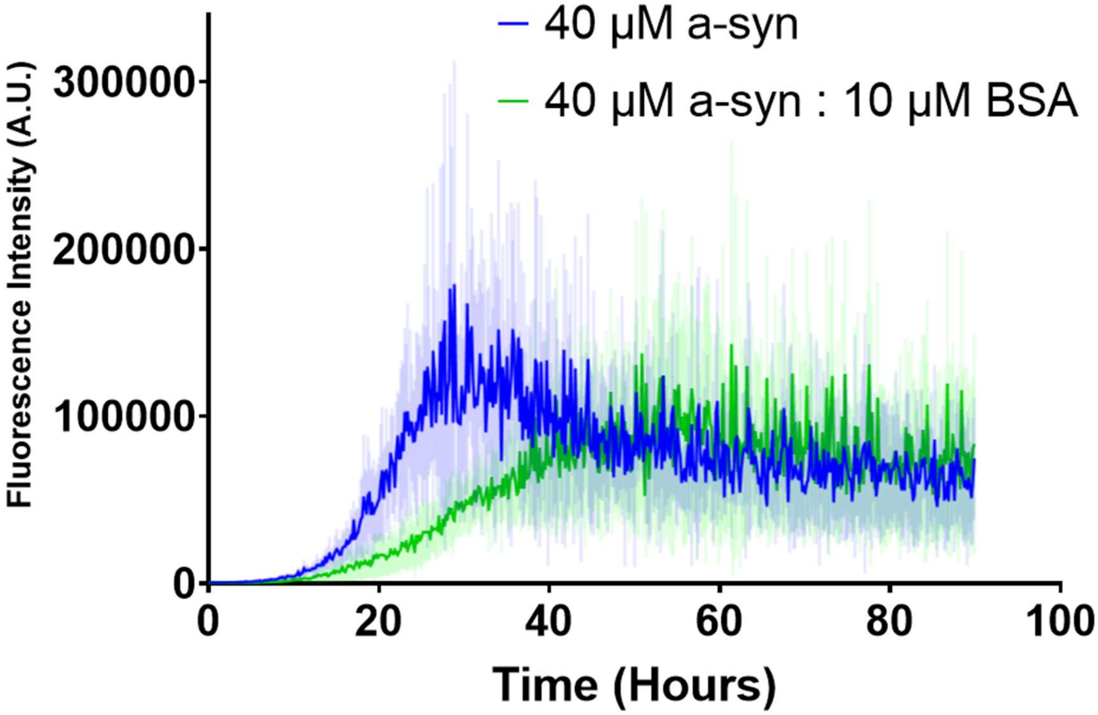


**Figure S5. The addition of BSA does not significantly reduce ThT fluorescence of a-Syn aggregation samples.** Aggregation kinetics of a-Syn (40 µM, 0.58 mg/ml) measured by ThT fluorescence in the absence (blue) or presence of Bovine serum albumin (BSA) (green; 10 µM, 0.67 mg/ml) at a similar mg/ml concentration equivalent to hTPR2A at 40 µM.

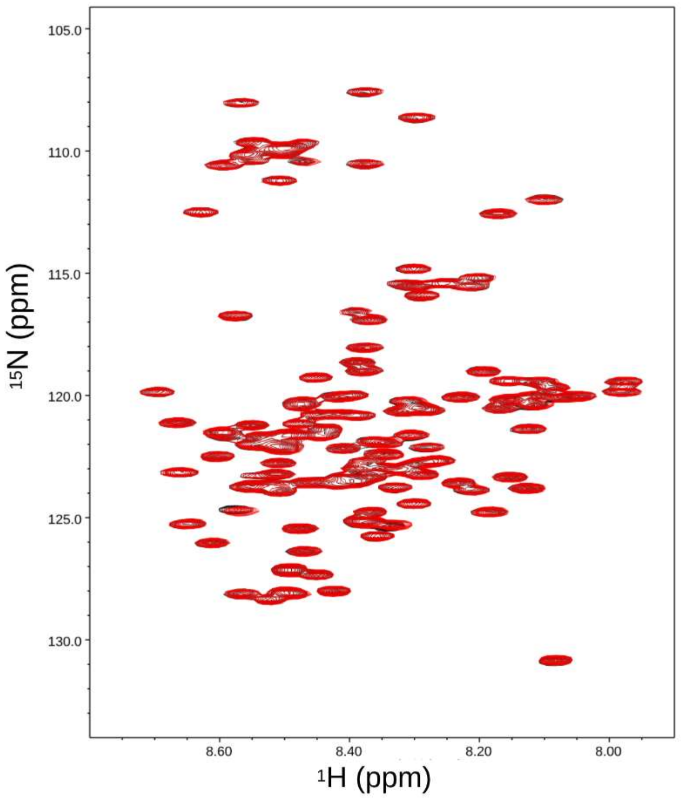


**Figure S6. Addition of TPR1 domain of STIP1 does not cause notable chemical shift changes of a-Syn.** $^{1}$H-$^{15}$N HSQC spectrum of 100 µM a-Syn in the absence (black) and the presence (red) of 200 µM of mTPR1.

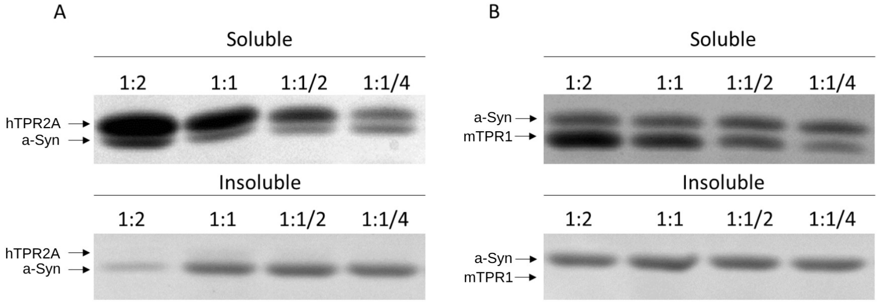


**Figure S7. The TPR2A domain of HOP maintains the solubility of a-Syn during aggregation.** Aggregation reactions containing 60 µM a-Syn in the presence of various concentration ratios of hTPR2A (A) or the non-specific binding control of mTPR1 (B) were incubated for 90 hours and separated into soluble and insoluble fractions by centrifugation.

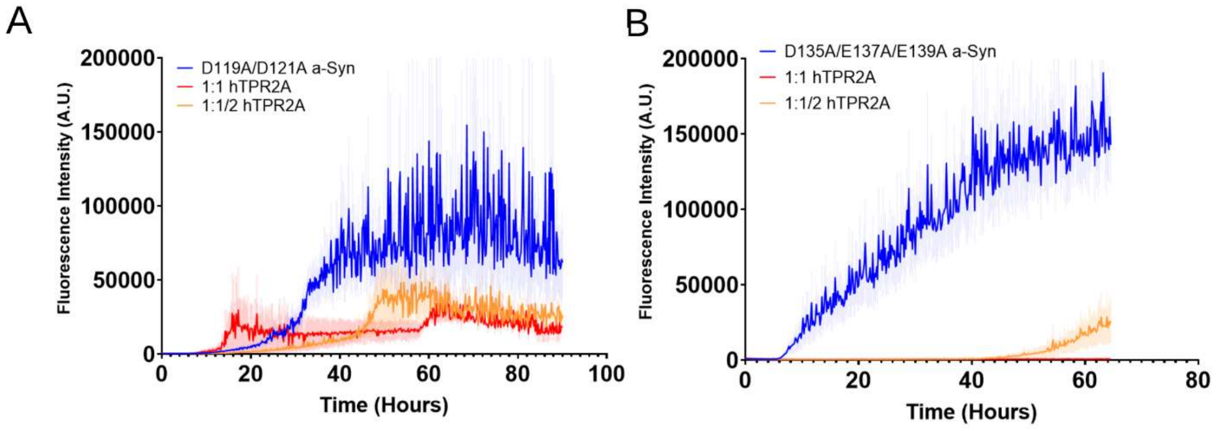


**Figure S8. hTPR2A inhibits fibril formation of D119A/D121A and D135A/E137A/E139A a-Syn.** (A) Fibril formation of D119A/D121A a-Syn (60 µM) measured by ThT fluorescence in the absence (blue) or presence of either equimolar (red) or half equimolar (orange) concentration of hTPR2A. (B) Fibril formation of D135A/E137A/E139A a-Syn (60 µM) measured by ThT fluorescence in the absence (blue) or presence of either equimolar (red) or half equimolar (orange) concentration of human hTPR2A.

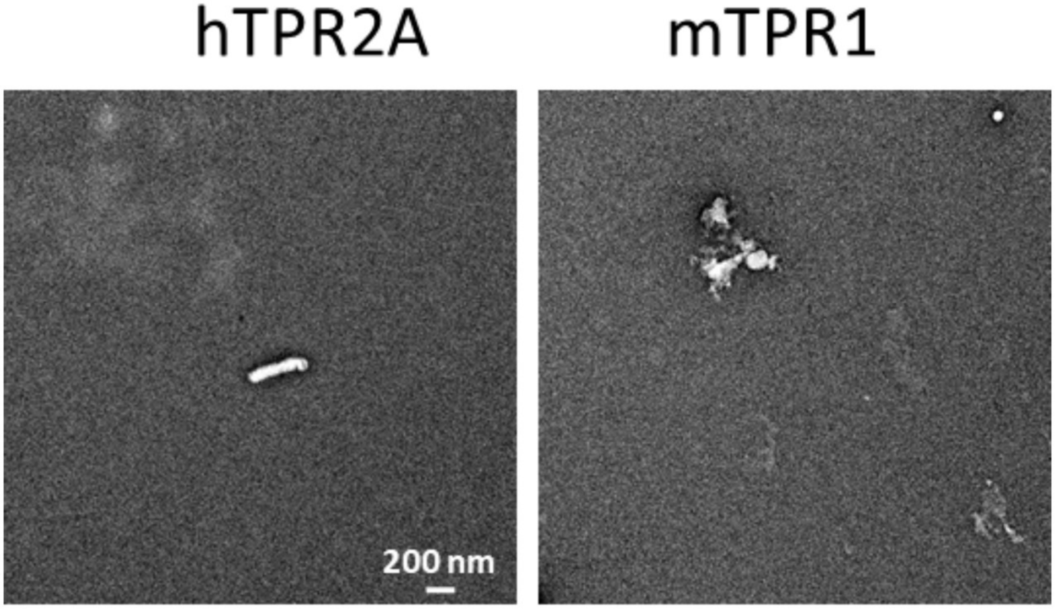


**Figure S9. Electron microscopy images of aggregated hTPR2A and mTPR1 control samples.** Representative EM images of precipitated species when present after 5 days for samples containing hTPR2A or mTPR1 undergoing standard a-Syn aggregation conditions in the absence of a-Syn.

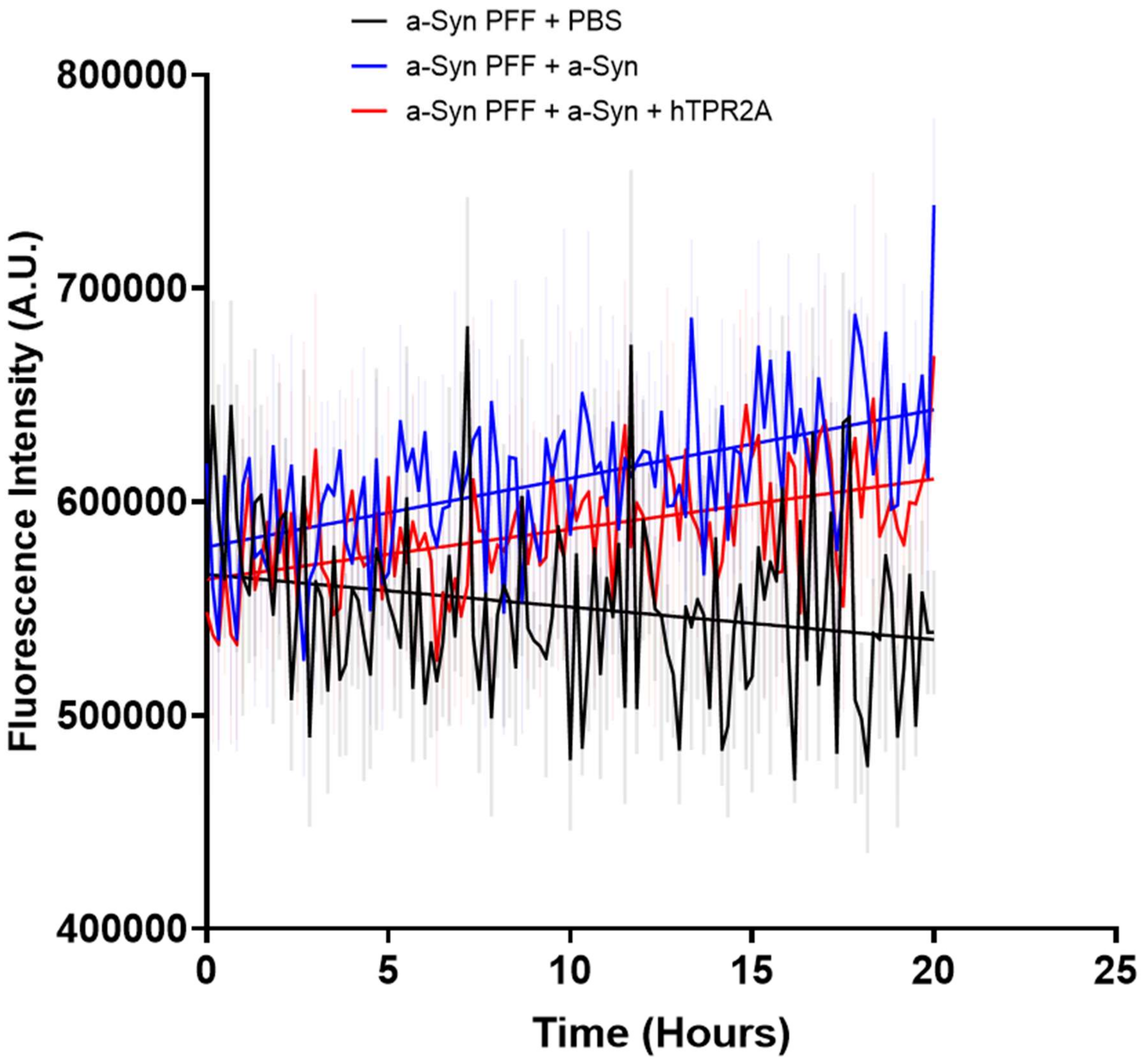


**Figure S10. hTPR2A does not prevent the incorporation of a-Syn to existing a-Syn fibrils.** Aggregation reactions containing 60 μM a-Syn were incubated for 90 hours, at which point ThT fluorescence plateaus. PBS buffer, a-Syn, or a-Syn with equimolar hTPR2A was then added and aggregation was continued.

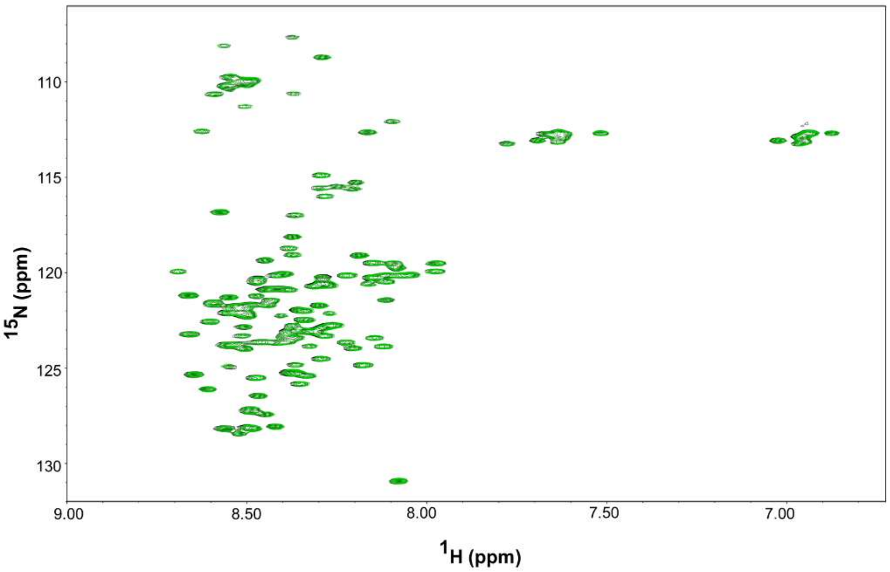


**Figure S11. The Hsp90 pentapeptide does not interact with a-Syn.** $^{1}$H-$^{15}$N HSQC spectrum of 100 μM a-Syn in the absence (black) and the presence (green) of 300 μM Hsp90 pentapeptide.

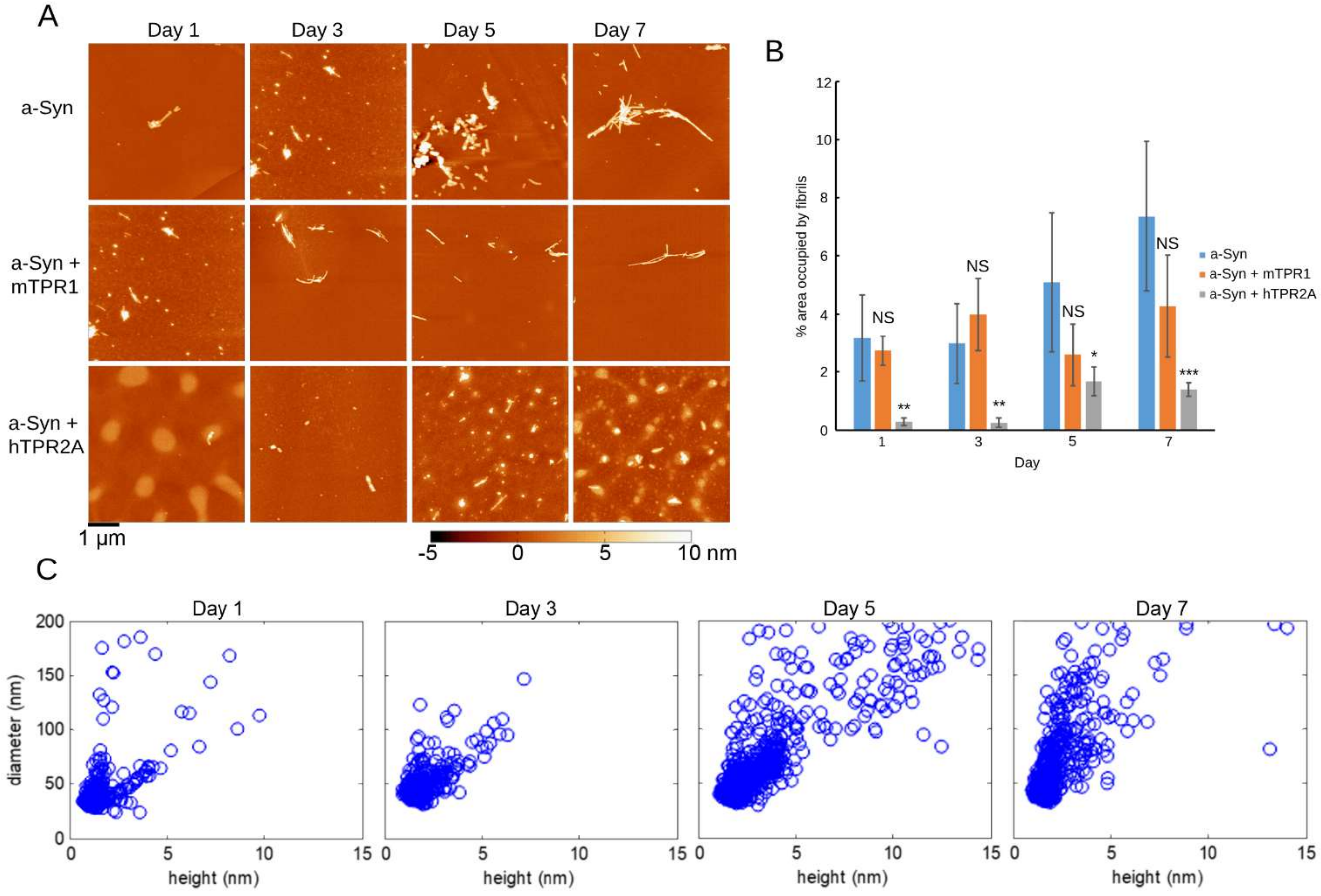


**Figure S12. Analysis of a-Syn assemblies formed in the presence of hTPR2A by AFM.** (A) Representative AFM images of a-Syn (175 µM) aggregation in the absence and presence of mTPR1 and hTPR2A at an equimolar concentration as a function of time. Protein samples are deposited on a mica surface. (B) Fibril load of a-Syn samples aggregated in the absence or presence of mTPR1 and hTPR2A as a function of time as estimated by area of AFM images occupied by fibrils. Statistical significance is represented by an asterisk, where **** is P < .0001, *** is P < .001, ** is P < .01, and * is P < .05. Error bars represent the standard deviation across a minimum of 5 images for each condition. (C) Correlation plots of the distribution of the heights and diameters of the amorphous high molecular weight a-Syn species produced in the presence of hTPR2A as a function of time.

**Movie S1.** Hierarchical cluster-ordered trajectory of a-Syn C-terminus binding to hTPR2A. Frames are arranged such that adjacent frames are most similar to each other. S1 motif residues are colored blue and S2 motif residues are colored yellow.
